# Multidirectional effect of low-intensity neuromuscular electrical stimulation on gene expression and phenotype in thigh and calf muscles after one week of disuse

**DOI:** 10.1101/2024.09.18.613609

**Authors:** Anna A. Borzykh, Roman Y. Zhedyaev, Ivan I. Ponomarev, Tatiana F. Vepkhvadze, Viktor G. Zgoda, Mira A. Orlova, Nikita E. Vavilov, Nikita V. Shishkin, Egor M. Lednev, Pavel A. Makhnovskii, Kristina A. Sharlo, Anastasia R. Babkova, Galina Yu. Vassilieva, Rinat R. Gimadiev, Boris S. Shenkman, Ilya V. Rukavishnikov, Oleg I. Orlov, Elena S. Tomilovskaya, Daniil V. Popov

## Abstract

**Purpose:** This study investigated the effects of a one-week disuse, both with and without low-intensity neuromuscular electrical stimulation – a safe (non-traumatic) approach to prevent the loss of muscle mass, on the functional capacities and gene expression in thigh and calf muscles.

**Methods:** This study assessed the efficiency of low-intensity (∼10% of maximal voluntary contraction) electrical stimulation in preventing the negative effects of 7-day disuse (dry immersion without and with daily stimulation) on the strength and aerobic performance of the ankle plantar flexors and knee extensors, mitochondrial function in permeabilized muscle fibers, and the proteomic (quantitative mass spectrometry-based analysis) and transcriptomic (RNA-sequencing) profiles of the soleus muscle and vastus lateralis muscle.

**Results:** Application of electrical stimulation during dry immersion prevented a decrease in the maximal strength and a slight reduction in aerobic performance of the knee extensors, as well as a decrease in maximal (intrinsic) ADP-stimulated mitochondrial respiration and changes in the expression of genes encoding mitochondrial, extracellular matrix, and membrane proteins in the vastus lateralis muscle. In contrast, for the ankle plantar flexors/soleus muscle, electrical stimulation had a positive effect only on maximal mitochondrial respiration, but slightly accelerated the decline in the maximal strength and muscle fiber cross-sectional area, which appears to be related to the activation of inflammatory genes.

**Conclusion:** The data obtained open up broad prospects for the use of low-intensity electrical stimulation to prevent the negative effects of disuse for “mixed” muscles, meanwhile, the optimization of the stimulation protocol is required for “slow” muscles.

**Practitioner Points:**

- Low-intensity electrical myostimulation is often used as an alternative to exercise and high-intensity electrical stimulation to prevent the loss of muscle mass and function in patients with severe chronic diseases and in spaceflight. However, its effect on muscles with different functional capacities remains uncertain.
- One week of disuse (dry immersion) lead to a comparable decrease in the maximal strength and (intrinsic) mitochondrial respiration in both the ankle plantar flexors/soleus muscle and the knee extensors/vastus lateralis muscle. Meanwhile changes in gene expression (transcriptome) were three times more pronounced in the soleus muscle than in the vastus lateralis muscle.
- Application of electrical stimulation during disuse prevented most of the negative effects of disuse in the knee extensors/vastus lateralis muscle, but accelerated the decline in the maximal strength/muscle fiber cross-sectional area in the ankle plantar flexors/soleus muscle, which may be related to the activation of genes regulating the inflammatory response.

## Introduction

The initial weeks of disuse (e.g., leg immobilization, bedrest, dry immersion), as well as spaceflight, cause a decrease in leg muscle strength and mass (Narici and de Boer 2011; Hackney and Ploutz-Snyder 2012; Koryak 2020; Vikne et al. 2020; Marusic et al. 2021) at a rate of 0.4%–0.5% per day (Narici and de Boer 2011; Hackney and Ploutz-Snyder 2012). Additionally, disuse leads to a reduction in the aerobic performance of the organism, measured as the maximum oxygen consumption rate in an incremental test on a cycle ergometer or treadmill, at a rate of 0.4%–1% per day (Ade et al. 2015; Ried-Larsen et al. 2017). Whole-body maximal aerobic performance is influenced by “central” factors (cardiac output) and “peripheral” factors (muscle O_2_ diffusion and oxidative capacity). However, only a few studies have investigated the effect of disuse on aerobic performance using exercise tests that involve small muscle mass – an approach eliminating the influence of “central” factors – such as tests focused solely on the knee extensors (Ringholm et al. 2011; Salvadego et al. 2011; Salvadego et al. 2016; Salvadego et al. 2018; Zuccarelli et al. 2021).

Low-intensity (<30% of maximal voluntary contraction) neuromuscular electrical stimulation (NMES) was offered as a safe (non-traumatic) approach and alternative to exercise and high-intensity NMES to prevent the loss of muscle mass, strength, and endurance in patients with severe chronic diseases (Quittan et al. 2001; Harris et al. 2003; Zanotti et al. 2003; Nuhr et al. 2004; Karavidas et al. 2010), as well as in spaceflight (Mayr et al. 1999; Shiba et al. 2015; Yarmanova et al. 2015; Maffiuletti et al. 2019). Interestingly, some effects of disuse, especially during dry immersion and spaceflight, manifest differently in leg muscles with different functional capacities. For instance, the soleus muscle – a muscle with strong postural function, which consists predominantly of type I slow muscle fibers, shows more pronounced effects than other leg muscles in terms of muscle strength, tone and mass, and muscle fiber size (LeBlanc et al. 1992; Kozlovskaya et al. 2007; Fitts et al. 2010; Tomilovskaya et al. 2019; Casuso et al. 2024). However, the effects of low-intensity NMES on strength and aerobic performance in various leg muscles are poorly investigated. This study assessed, for the first time, the efficiency of low-intensity NMES in preventing the negative effects of strict disuse on both the ankle plantar flexors/soleus muscle and the knee extensors/vastus lateralis muscle, using a dry immersion model. Specifically, we investigated the effects of a one-week dry immersion, both with and without NMES, on the strength and aerobic performance of leg muscles (test involving small muscle mass), mitochondrial function in permeabilized muscle fibers, and the proteomic (quantitative mass spectrometry-based analysis) and transcriptomic (RNA-sequencing) profiles of skeletal muscles.

## Methods

### Ethical approval

The studies were approved by the Committee on Biomedical Ethics of the Institute of Biomedical Problems of the RAS (protocols № 594 of September 6, 2021, and № 620 of July 12, 2022) and complied with all the principles set forth in the Declaration of Helsinki. All volunteers gave written informed consent to participate in the study and to undergo skeletal muscle biopsy.

### Study design

Ten males participated in a control 7-day dry immersion (DI, ages 25–38 years; median height 1.78 m [interquartile range 1.71–1.79 m]; body mass 70 kg [64–74 kg]; and body mass index 24 kg/m² [21–24 kg/m²]). A year later (at the same time of year), another ten males participated in a 7-day dry immersion with NMES (DI+NMES, ages 26–39 years; height 1.74 m [1.70–1.76 m]; body mass 73 kg [70–79 kg]; and body mass index 24 kg/m² [23–26 kg/m²]). Study design and test protocols were similar in both experiments.

To evaluate the effects of NMES, changes induced by DI were compared with changes induced by DI+NMES. After a familiarization session, one day before and 8–9 hours after the end of DI or DI+NMES, the maximum isometric voluntary contraction (MVC) of the ankle plantar flexors and knee extensors of the right leg and (after 10 minutes of rest) the aerobic performance of these muscles (maximum power in an incremental ramp test till exhaustion, *W_max_*) were examined (Fig. 1A). To evaluate markers of NMES-induced muscle damage the venous blood was drawn from the *v. intermedia cubiti* 7 days before and 4 days after the start of disuse. Muscle samples were taken using a Bergstrom needle with aspiration under local anesthesia (2 ml of 2% lidocaine) from the medial part of the vastus lateralis (VL) and soleus (Sol) muscles of the left leg 14 days before and 6 days after the start of disuse (DI and DI+NMES) at 10:00, 3 hours after a standardized very light breakfast (5.2 g protein, 2.7 g fat, 55 g carbohydrates, 1253 kJ). This biopsy time was chosen because various physiological tests were performed immediately after the end of the 7-day dry immersion (not included in this study); this approach allowed us to study the “pure” molecular response to 6 days of disuse, as well as the “relatively pure” physiological response to 7 days of disuse. The samples were cleared of visible fragments of connective and adipose tissue; part of the fresh tissue was used to assess mitochondrial function, another part was immediately frozen in liquid nitrogen to study the transcriptome and proteome profiles, and the last part was embedded in Tissue-Tek O.C.T. Compound (Sakura, USA) and frozen in liquid nitrogen to histological study; all frozen samples were stored at -80 °C until further analyses.

**Fig. 1.**
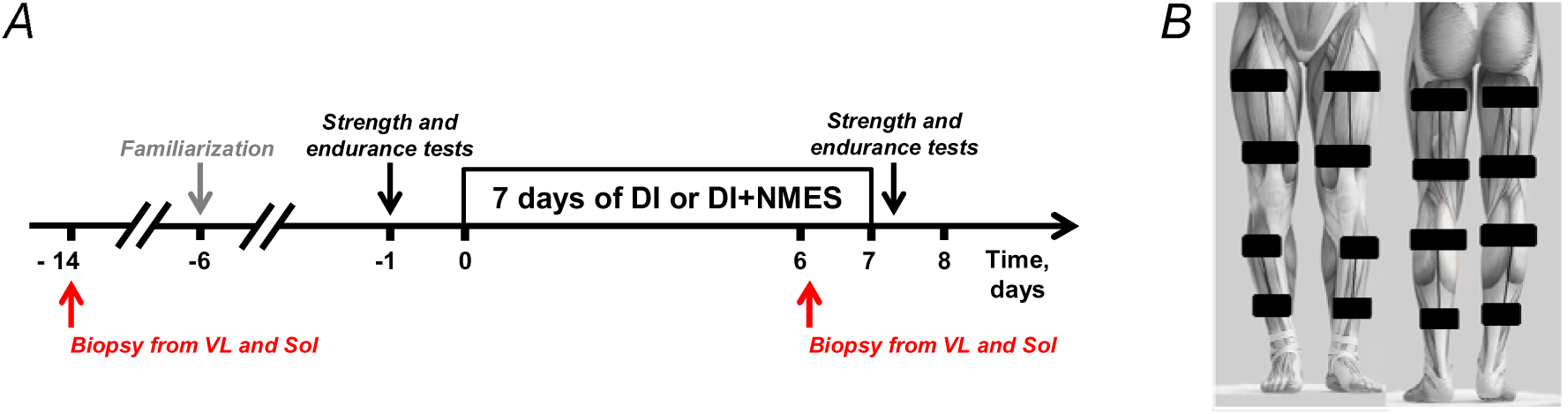
Design of the study. A – Dry immersion without (DI) and with neuromuscular electrical stimulation (DI+NMES). B – Position of electrodes for NMES.

### Dry immersion protocol

In both experiments during dry immersion subjects remained immersed in a bath with thermoneutral water (32.7±0.7 degrees C) in a supine position for all activities. Light-off period was set at 23:00-07:00. Daily hygiene and some measurements (not included in the paper) required to pull a person out from the bath. Average out-of-bath time was: 19.6±5.7 min (including 7.0±3.9 min in vertical position) on days 1-5; 51.5±8.8 min (including 7.5±1.7 min in vertical position) on day 6, and 73.5±7.0 min (including 5.9±1.7 min in vertical position), because of muscle biopsy, on day 7. Blood pressure, heart rate, and body temperature were measured three times per day (8:00, 15:00, and 21:00); subjects were constantly monitored with video. Dietary intake was controlled during the study; the menu composition was identical for all subjects.

### Neuromuscular electrical stimulation

Motor neuron firing rate plays an important role in the activation of calcium-dependent signaling and the expression of muscle-specific genes. Therefore, in our study, both low (25 Hz) and high (50 Hz) stimulation frequencies were used during DI+NMES (in the first and second NMES daily session, respectively).

Every day at 11:00 (7 sessions in total), low-intensity low-frequency electrical stimulation was applied to the calf and thigh muscles for 40–45 min (bipolar symmetrical rectangular pulses [1 ms, 25 Hz] for 1 s, followed by a 2-s pause) using a stimulator Stimul-01 (Biophyspribor, Russia), as described elsewhere (Poltavskaya et al. 2021).

Every day at 17:00 (7 sessions in total), low-intensity high-frequency electrical stimulation was applied to the calf and thigh muscles for 10 min (carrier frequency of 2-kHz bipolar symmetrical pulses [10 ms separated by 10-ms pause] with bursts of 50 Hz for 10 s, followed by a 50-s pause) using a stimulator Amplidin-EST (NTC UP RAN, Russia), as described elsewhere (Amirova et al. 2022). The burst time during high-frequency NMES was longer than that during low-frequency NMES (10 s *vs.* 1 s). Therefore, to increase the pain tolerance during high-frequency NMES, the carrier frequency (2-kHz) was used (Ward and Shkuratova 2002).

Electrodes (13 cm × 5 cm, PG-901, Fiab, Italy) were placed on the proximal and distal thirds of the anterior and posterior surface of the thigh. For the posterior calf muscles, one electrode (13 cm × 5 cm) was placed in the proximal part of the calf, and another electrode (9 cm × 5 cm) – 5 cm distal to the gastrocnemius muscles. For the anterior calf muscles, electrodes (9 cm × 5 cm) were placed on the proximal and distal thirds of the anterior surface of the calf (Fig. 1B). NMES was performed in a bath: the subject was slightly raised to the surface of the water using the bath lifting platform and lay in a supine position: the knee angle joint was 160-165 degrees (180 degree – knee fully extended), the angle at the ankle joint was 110 degrees or 130 degrees during NMES-induced contraction of the calf muscles (90 degree – foot perpendicular to the tibia, greater values represent plantar flexion angles).

This study focused on investigating the effects of low-intensity NMES, which does not cause significant unpleasant sensations and, therefore, can potentially be used for patients with various diseases or healthy people to countermeasure negative effect of strict disuse. Therefore, during DI+NMES, the intensity of stimulation was individually adjusted to the point of causing unpleasant sensations. The mean voltage [standard deviation] of low-frequency stimulation was 19.2 ± 4.2 V for the anterior thigh, 17.7 ± 4.6 V for the anterior calf, 17.6 ± 4.2 V for the posterior thigh, and 18.2 ± 4.3 V for the posterior calf. The mean current during high-frequency stimulation was: for both anterior and posterior surfaces of the thigh – 18.3 ± 5.2 mA and calf – 10.5 ± 2.5 mA. NMES caused visible contractions of the stimulated muscles (∼10% MVC for both knee extensors and ankle plantar flexors). Namely, the NMES-induced relative force of the knee extensors (or ankle plantar flexors) at the stimulation intensity assessed as “the point of causing unpleasant sensations” was examined in an additional experiment. Each subject involved in the DI experiment sat on a Pro System 3 dynamometer (Biodex, USA): the knee joint was 135 degrees (180 degree – knee fully extended). To investigate the NMES-induced relative force of the ankle plantar flexors the subjects were lying in a supine position with one leg fixed on a homemade isometric ankle plantar flexors dynamometer: the ankle joint was 90 degrees (foot perpendicular to the tibia) and the knee joint was 135 degrees. The stimulation intensity of the knee extensors (or ankle plantar flexors) was gradually increased and the force (torque) was recorded, as well as “the point of causing unpleasant sensations”. Then, the isometric MVC at this position was examined (Supporting information Fig. S1), as described in the next section.

### Maximum voluntary contraction of the ankle plantar flexors and knee extensors

Isometric MVC of plantar flexors was assessed in a sitting position (Supporting information Fig. S2). To reduce the contribution of the biarticular gastrocnemius muscles to force production (Cresswell et al. 1995; Niess et al. 2018), MVC of the ankle plantar flexors was examined at a knee joint angle of 70 degrees, the upright position of the calf, and the ankle joint 90 degrees using a CalfRaise-PRO dynamometer (AntexLab, Russia). A force transducer was pressed down to the knee joint (200 N) and then reset to zero.

The isometric MVC of the knee extensors was examined in a sitting position on a Pro System 3 dynamometer (Biodex, USA) at a knee joint angle of 110 degrees.

In each test, the subject performed 5–7 attempts with a rest period of at least 30 s between attempts; the best attempt was taken into account. Force/torque analog signal from both dynamometers was recorded using an analog-to-digital converter E14-440 (L-CARD, Russia) and PowerGraph 3 software (Interoptica-c, Russia) with sampling frequency 1 kHz (Supporting information Fig. S2 and S3).

### Maximum aerobic performance of the ankle plantar flexors and knee extensors

Maximum aerobic power (*W_max_*) of the ankle plantar flexors was assessed in an incremental ramp test to exhaustion in a sitting position, as described above. Additionally, the distal part of the foot was placed on a 4 cm high stand (Supporting information Fig. S2); the subject performed rhythmic extensions of the metatarsophalangeal joint (raising the calf with 55% of the maximum amplitude): raising – 1.1 s, lowering – 0.3 s, rest – 0.6 s. The rhythm and amplitude of movement were set using biofeedback (analog-to-digital converter E14-440 and PowerGraph 3 software). The incremental test was carried out using the CalfRaise-PRO ergometer: the initial load for raising the calf was 30 N, the load increment was 25 N/min, and the load during calf lowering was minimal. The test was stopped when the subject was unable to maintain the full amplitude at the given rhythm for five cycles. The power for each cycle was calculated as

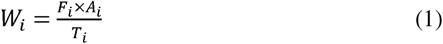

where *i* – cycle number, *F_i_* – average load during muscle contraction in a cycle (N); *A_i_* – amplitude of movement in a cycle (m); *T_i_* – cycle duration (s). The dependence “*W_i_* – time” was smoothed (moving average with a window of 20 s), then *W_max_* was defined as the maximum value (Supporting information Fig. S2).

The maximum aerobic power of the knee extensors was assessed using a modified Ergoselect 900 cycle ergometer (Ergoline, Germany): each subject performed rhythmic knee extensions (from 80 to 160 degrees) in a reclining sitting position (hip joint 35 degrees) (Supporting information Fig. S3). The extension frequency (1 cycle/s) was set by a sound command, the initial load – 0 W, the load increment – 5 W/min, and the load during flexion was minimal. During the test, constant cadence (1 cycle/s) was simulated on the ergometer cadence transducer. The amplitude of knee extension was ∼80 degrees, but the frequency of extension could be different from the specified (simulated) one (because of fatigue), therefore the power per cycle was calculated as

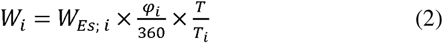

where *i* – cycle number, *W_Es; i_* – power per cycle set on the Ergoselect 900 ergometer (W); φ_i_ – change in the knee joint angle per cycle (degrees); *T* and *T_i_*– specified and actual cycle duration, respectively (s). The knee joint angle was recorded using the analog-to-digital converter E14-440 and PowerGraph 3 software. The test was stopped when the subject was unable to maintain the full amplitude in the given rhythm for 5 cycles. The dependence “*W_i_* – time” was smoothed (moving average with a window of 20 s), and then *W_max_* was defined as the maximum value (Supporting information Fig. S3).

### Serum creatine kinase (CK) activity and myoglobin level

CK activity was measured by an automatic analyzer Cobas c502, and myoglobin – a Cobas e601 (both Roche Diagnostics, Germany) using standard reagents.

### Muscle fiber cross-sectional area (CSA)

Muscle transverse sections (10 μm) were prepared using a cryotome (CM 1900, Leica Biosystems, Germany), and immunostaining was performed using primary antibodies for slow (protein product of *MYH7*, 1:100, M8421, RRID:AB_477248, Sigma, USA) and fast muscle fibers (protein product of both *MYH1* and *MYH2*, 1:100, SC-75/ACC 232, RRID:AB_3106894, Leibniz Institute DSMZ, Germany), as well as secondary antibodies AlexaFluor 488 and 546 (A32723, RRID:AB_2633275 and A11035 RRID:AB_2534093), respectively, all 1:400, Thermo Fisher Scientific, USA). Images were acquired using a fluorescent microscope Olympus IX83P2ZF3 (Olympus, Japan; 20× objective magnification) and analyzed by the ImageJ software. The median (range) number of muscle fibers assessed was 101(54 – 121) for VL in DI+NMES, 89(20 – 134) for Sol in DI+NMES, 76(54 – 132) for VL in DI, and 59(20 – 101) for Sol in DI.

### Mitochondrial respiration in permeabilized muscle fibers

A piece of fresh tissue (∼10 mg) was placed in ice-cold BIOPS relaxation buffer (10 mM Ca-EGTA buffer, 0.1 μM free calcium, 20 mM imidazole, 20 mM taurine, 50 mM K-MES, 0.5 mM DTT, 6.56 mM MgCl_2_, 5.77 mM ATP, 15 mM phosphocreatinine, pH 7.1) to remove connective and adipose tissue. Then a small piece (∼2 mg) was removed and the muscle fibers were separated using a pair of needles to form a bundle (two bundles for each biopsy sample). The fibers were permeabilized in BIOPS buffer with saponin (50 μg/ml, 30 min) on ice with slow stirring, then the tissue was washed 2 times for 10 min in MiRO5 buffer (0.5 mM EGTA, 3 mM MgCl_2_·6 H_2_O, 60 mM lactobionic acid, 20 mM taurine, 10 mM KH_2_PO_4_, 20 mM HEPES, 110 mM D-sucrose, 1 g/l fatty acid-free bovine serum albumin). The rate of mitochondrial respiration was assessed simultaneously on two channels on an Oxygraph polarograph (Hansatech, United Kingdom) under hyperoxic conditions (Co_2_ >200 μM) to avoid potential oxygen limitation. Nonphosphorylating leak respiration was assessed by sequentially adding: malate (2 mM) + pyruvate (5 mM) + glutamate (10 mM). Maximal coupled (ADP-stimulated) respiration with convergent electron input to complexes I and II of the electron transport system was achieved by adding ADP (5 mM) + MgCl_2_ (3 mM), followed by succinate (10 mM). The integrity of the outer mitochondrial membrane was tested by adding cytochrome *c* (10 μM; if a respiration increase <10%, the quality of the mitochondria was considered sufficient). Then the FCCP uncoupler was titrated (0.25 μM to 1.5 μM) to estimate the maximum capacity of the electron transport chain (maximum uncoupled respiration). The respiration rate was normalized to the mass of permeabilized muscle fibers, then, the average respiration rate for two channels was calculated.

### RNA sequencing and data processing

A piece of frozen tissue (∼15 mg) was homogenized at +4 °C in 500 μl of a lysis buffer (ExtractRNA, Evrogen, Russia) using a plastic pestle and a drill. Total RNA was isolated using a spin column (RNeasy Mini Kit, Qiagen, Germany). RNA concentration was assessed using a Qubit 4 fluorimeter (ThermoScientific, USA); RNA integrity – using capillary gel electrophoresis (TapeStation, Agilent, USA). All samples had RNA integrity (RIN) >7.8.

Strand-specific RNA libraries were prepared, as described previously (Kurochkina et al. 2024), using the NEBNext Ultra II RNA kit (New England Biolabs, USA) according to the manufacturer’s recommendations. Library efficient concentration was evaluated by qPCR using 5X qPCRmix-HS SYBR kit (Evrogen, Russia); single-end sequencing was done by a NextSeq 550 analyzer (Illumina, USA), according to the manufacturer’s recommendations, with a read length of 75 bp (∼60 million reads/sample).

Sequencing quality was assessed using the FastQC tool (v.0.11.5, RRID:SCR_014583), and low-quality reads were removed using Trimmomatic (v.0.36, RRID:SCR_011848). High-quality reads were mapped to the human reference genome GRCh38.p13 primary assembly. The number of unique reads mapped to exons of each gene was determined using the Rsubread package (R environment) and Ensembl annotation (GRCh38.101). The DESeq2 method (paired sample analysis with Benjamini-Hochberg correction) was used to analyze differential gene expression. Differentially expressed genes were defined as protein-coding genes (as well as polymorphic pseudogenes and translated pseudogenes) with an expression level of TPM >1 (Transcripts Per Million, kallisto tool v0.46.2), *p_adj_*<0.01 and |fold change|≥1.25. A comparison of VL before DI *vs.* Sol before DI, as well as each muscle after DI (or DI+NMES) *vs.* the same muscle before DI (or DI+NMES) was performed.

### Quantitative mass spectrometry-based proteomic analysis

A piece of frozen tissue (∼10 mg) was homogenized as described above in 140 μl of lysis buffer (4% sodium dodecyl sulfate, 0.1 M Tris and 0.1 M dithiothreitol, pH 7.6). The lysate was boiled (95 °C, 5 min), transferred to an AFA microtube and sonicated (average power 20 W, 30 s × 4 using a ME220 sonicator (Covaris, USA), and centrifuged (5 min, 30,000 g). The protein concentration in the supernatant was measured by a fluorimeter Qubit 4.

The proteins (85 μg) were diluted in lysis buffer (final volume 24 μl), transferred to an S-Trap column (ProtiFi, USA), alkylated (20 mM iodoacetamide, 15 min), and hydrolyzed into the S-Trap column using trypsin and Lys-C (2 h at 47 °C, 1:15 and 1:30, respectively, all Promega, USA) in 40 μl 50 mM triethylammonium bicarbonate according to the manufacturer’s recommendations. After elution, 20 μg of peptides (fluorimeter Qubit 4) were dried, resuspended in 100 mM triethylammonium bicarbonate, and labeled with TMT 10-plex or 16-plex isobaric labels (Thermo Scientific, USA) for 1.5 h according to the manufacturer’s recommendations. The reaction was stopped (0.3% hydroxylamine w/v, 15 min), and samples were pooled.

The mixture of labeled peptides was separated using high pH reverse-phase LC fractionation (HPLC 1200, Agilent, USA). The peptides were concentrated on an analytical column (XBridge C18, particle size 5 μm, 4.6 x 250 mm, Waters, Ireland) in isocratic mode at a flow of 750 μl/min for 3 min in mobile phase A (15 mM ammonium acetate, pH 9,0). Twenty-four fractions were collected from 3 min to 50 min (collection time 2 min, volume 1500 μL) using a gradient elution mode with mobile phase B (15 mM ammonium acetate, pH 9.0, 80% acetonitrile, pH 9.0). Each fraction was concentrated, and then fractions (1 and 13, etc.) were combined to obtain 12 fractions.

Each fraction was analyzed three times using an HPLC Ultimate 3000 RSLC nanosystem (Thermo Scientific, USA) and a Q Exactive HF-X Hybrid Quadrupole-Orbitrap mass spectrometer (Thermo Scientific, USA) by the nanoelectrospray ion source in the positive mode of ionization (Thermo Scientific) as previously described (Kurochkina et al. 2024). The gradient (90 min) was formed by the mobile phase A (0.1% formic acid) and B (80% acetonitrile, 0.1% formic acid) at a 0.4 μL/min flow. The ionizing voltage was 2.1 kV. MS spectra were acquired at a resolution of 60,000 in the 390–1300 m/z range; fragment ions were mass scanned at a resolution of 60,000 at the range from m/z 120 to the upper m/z value as assigned by the mass to charge state of the precursor ion. All tandem MS scans were performed on ions with a charge state from z = 2+ to z = 4+. Synchronous precursor selection facilitated the simultaneous isolation of up to 40 MS2 fragment ions. The maximum ion accumulation times were set to 50 ms for precursor ions and 25 ms for fragment ions. AGC targets were set to 10^6^ and 10^5^ for precursor ions and fragment ions, respectively.

Search and identification of peptides and proteins were performed using a MaxQuant platform (2.1.4.0; Max Planck Institute for Biochemistry, RRID:SCR_014485) with default settings (FDR for peptides 1%, N-terminal acetylation and oxidation of methionine as variable modifications and carbamidomethylation of cysteine, as fixed modification) and the Isobaric much between runs and PSM-level weighted ratio normalization functions (Yu et al. 2020). Further analysis was performed using a Perseus platform (1.6.5; Max Planck Institute for Biochemistry, RRID:SCR_015753). To avoid the batch effect associated with labeling with the TMT 10-plex and 16-plex kits, the average reporter ion intensity of all samples labeled with 10-plex isobaric tags was normalized to the intensity of all samples labeled with 16-plex tags, as previously described (Kurochkina et al. 2024). After filtration (potential contaminants, reverse peptides, peptides identified by site only), proteins identified by more than one peptide (unique+razor) and present in >70% of samples were taken for further analysis. Differentially expressed proteins were determined using one-sample Wilcoxon signed tank test with *q*-value (Benjamini-Hochberg adjusted *p*-value) <0.05 and Tukey’s *post-hoc* test (*p*<0.05). A comparison of VL before DI *vs.* Sol before DI, and each muscle after DI *vs.* the same muscle before DI was performed.

### Western blot analysis

The lysate samples were mixed with Laemmli buffer and loaded onto 10% polyacrylamide gels (15 μg protein/lane), as described elsewhere (Popov et al. 2018). Electrophoresis (20 mA per gel) was performed using a Mini-PROTEAN Tetra Cell system (Bio-Rad, USA). The proteins were transferred to nitrocellulose membranes using a Trans-Blot Turbo system (Bio-Rad) in Towbin buffer for 30 min at 25 V. The membranes were stained with Ponceau S to verify consistent loading of proteins, followed by washing and incubation for 1 h in 5% non-fat dry milk (Applichem, Germany) in TBSt (20 mM Tris-HCl, pH 7.6; 150 mM NaCl, 0.1% Tween 20). Next, the membranes were incubated at 4°C overnight with antibodies specific oxidative phosphorylation proteins NDUFB8, SDHB, UQCRC2, COX1 and ATP5A1 (1:2000; ab110413; Abcam) in 5% non-fat dry milk in TBST. Blots were then incubated for 1 h at room temperature with an anti-mouse Ig secondary antibody (1:5000, Cell Signaling Technology) in 5% non-fat dry milk in TBST. After each step, the membranes were washed three times for 5 min with TBST. Finally, the membranes were incubated with SuperSignal West Femto Maximum Sensitivity Substrate (Thermo Scientific, USA) and luminescent signals were captured using a ChemiDoc Imaging System (Bio-Rad).

### Statistical analysis

Data are presented as median and interquartile range. A comparison of MVC, *W_max_*, fiber CSA, and mitochondrial respiration in each muscle group after DI (or DI+NMES) *vs.* the same muscle group before DI (or DI+NMES) was performed using the nonparametric Wilcoxon test (Fig. 2). A comparison of changes in each of these parameters caused by DI and DI+NMES was performed using the nonparametric Mann-Whitney test (Fig. 2). A comparison of mitochondrial respiration in VL before DI *vs.* Sol before DI was performed using the nonparametric Wilcoxon test (Fig. 3).

**Fig. 2.**
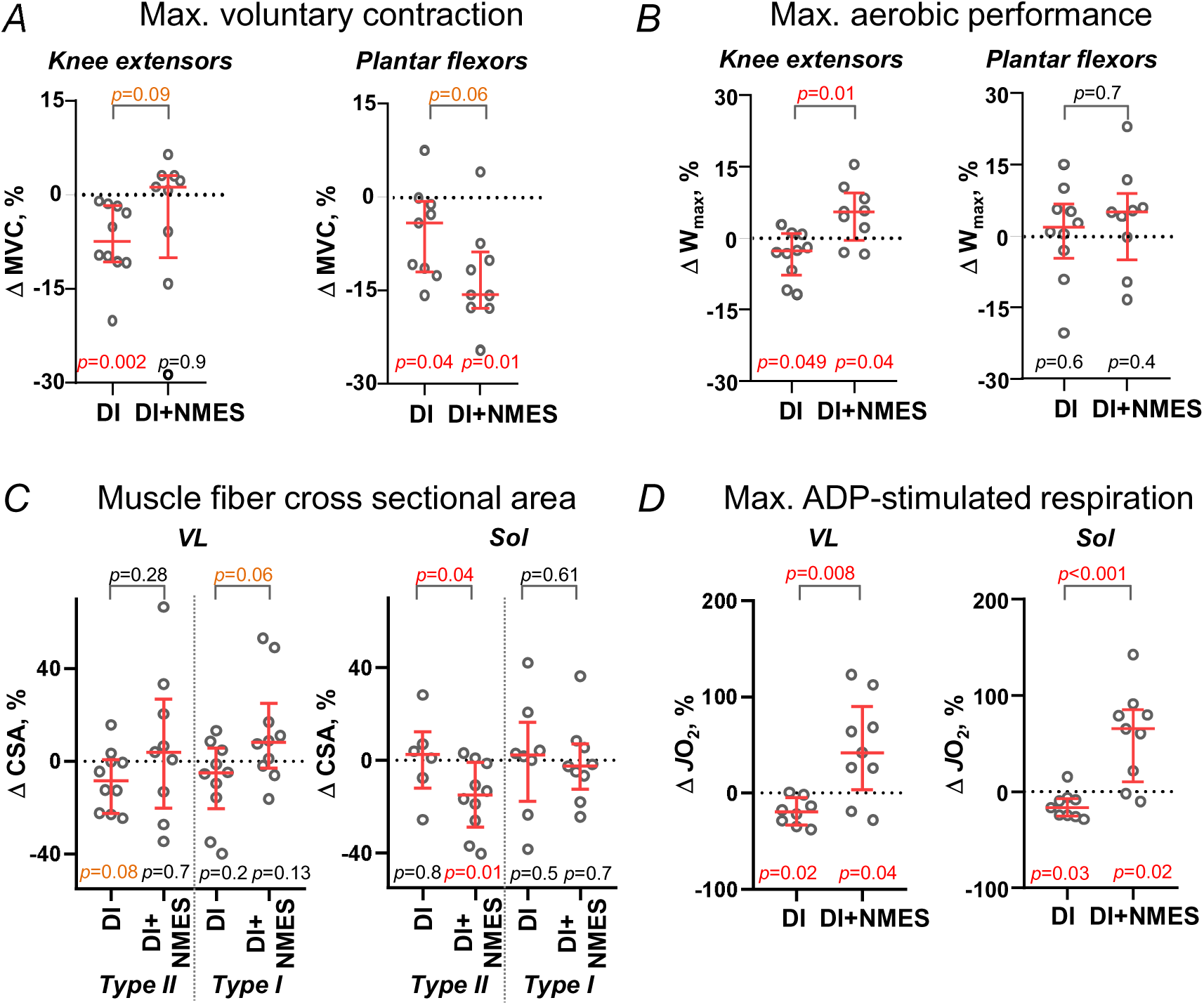
Electrical stimulation prevented the dry immersion-induced decrease in maximal voluntary contraction (MVC) and a slight decrease in the endurance of the knee extensors, but increased the drop in both MVC and type II muscle fibers cross-sectional area (CSA) in the ankle plantar flexors/soleus muscle. A and B – Changes in isometric maximum voluntary contraction (A) and aerobic performance (maximum power in the incremental ramp test, *W_max_*) (B) in the knee extensors and ankle plantar flexors after dry immersion without (DI) and with neuromuscular electrical stimulation (DI+NMES); (*n* = 9–10 subjects in each group). C and D – Changes in the cross-sectional area of type I and II muscle fibers (C, *n* = 6–10 in each group) and maximum ADP-stimulated mitochondrial respiration (*Jo_2_*) in permeabilized fibers (D) in VL and Sol muscles; (*n* = 9 subjects in each group). Median, interquartile range, and individual values are shown. The lower *p*-values refer to changes from the baseline (dashed line) (Wilcoxon signed-rank test), the upper ones refer to differences between the magnitude of changes in DI and DI+NMES (Mann-Whitney test).

**Fig. 3.**
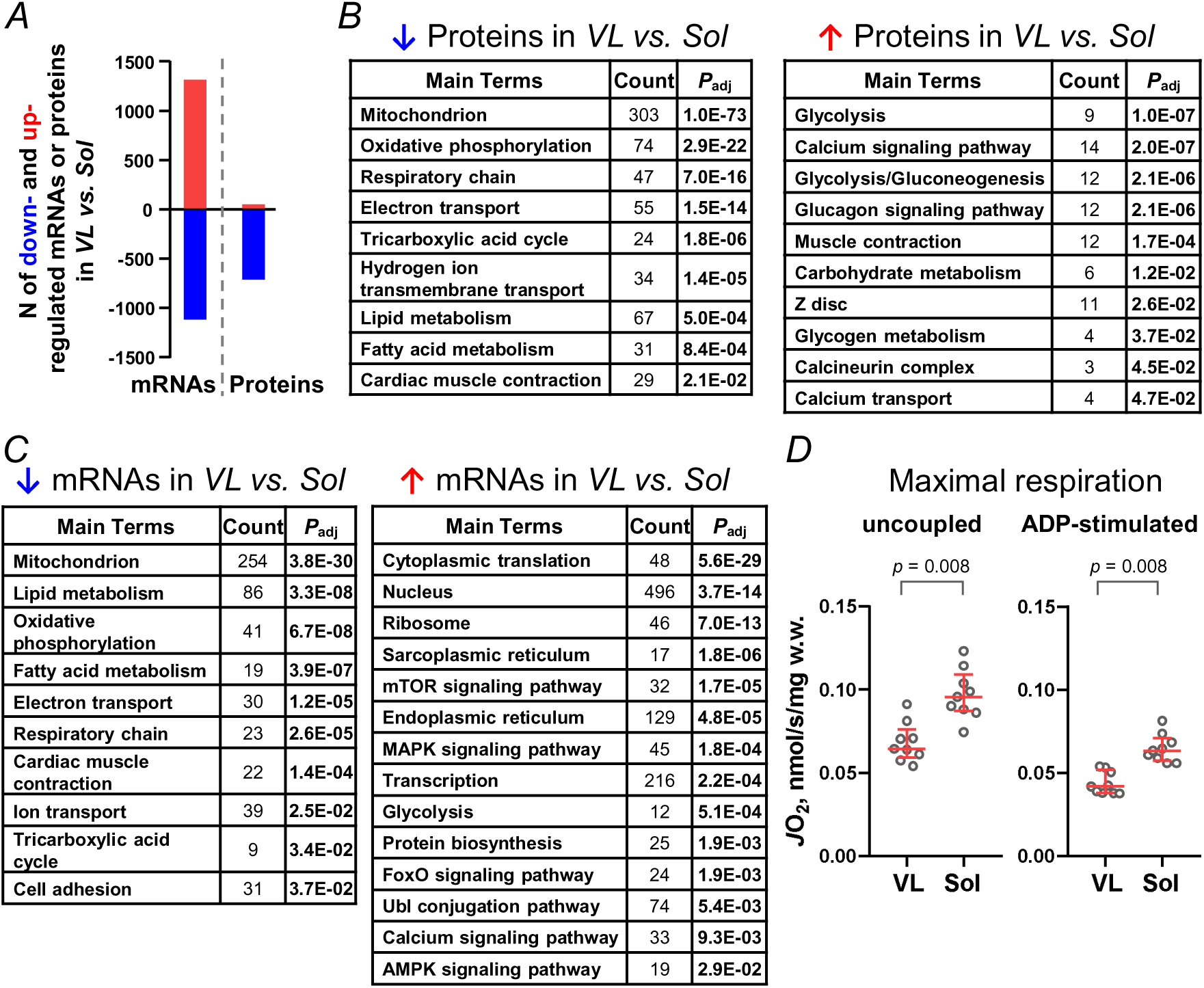
Pronounced differences between the vastus lateralis (VL) and soleus (Sol) muscles in the baseline (prior to dry immersion) for proteomic and transcriptomic profiles and mitochondrial respiration. A – number up- and down-regulated mRNA/proteins in VL *vs.* Sol in the baseline (*n* = 10 subjects in each group). All detected protein-coding genes and proteins are presented in Supporting information Table S1. B and C – Tables show the number of mRNA/proteins and the corresponding *p_adj_*for the main enriched functional categories (all results of enrichment analysis are presented in Supporting information Table S2). D – Maximum uncoupled and ADP-stimulated mitochondrial respiration (*Jo_2_*) in permeabilized fibers in VL and Sol in the baseline. Median, interquartile range, and individual values are shown. *p*-values refer to differences between the magnitude of change in DI and DI+NMES (Mann-Whitney test, *n* = 9 subjects in each group).

For transcriptomic and proteomic data, functional enrichment analysis for biological processes, cellular components and pathways (UNIPROT KW BP/CC, GENE ONTOLOGY BP/CC DIRECT and KEGG PATHWAY databases) was performed against all protein-coding genes (TPM >1, ∼10,000 mRNA) or detected proteins using the DAVID tool v2023q4 (RRID:SCR_001881) (*p_adj_*<0.05, Fisher’s exact test with Benjamini correction).

## Results

### Strength and endurance of the knee extensors and ankle plantar flexors

DI resulted in comparable reductions in the knee extensor and ankle plantar flexor MVC (-7(-11 – -2)% and -4(-12 – -1)%, respectively, *p*<0.05; Fig. 2A). NMES had different effects on different muscles: it prevented the dry immersion-induced decrease in MVC in the knee extensors (*p* = 0.9), but unexpectedly increased the drop in strength in the ankle plantar flexors (up to -16(-18 – -9)%, *p* = 0.01). Moreover, a trend toward a difference was found between the magnitude changes in these measures in DI and DI+NMES for both muscle groups (*p* = 0.06– 0.09), indicating the multidirectional effect of NMES on various muscle groups (Fig. 2A).

DI caused a very small decrease in aerobic performance in the knee extensors only (-3(-8 – 1)%, *p* = 0.049) (Fig. 2B). In contrast, DI+NMES led to an increase in this measure (by 5(0 – 10)%, *p* = 0.039); moreover, the magnitude of change in these measures after DI differed (*p* = 0.01) from that after DI+NMES, underlining the positive effect of NMES on VL. Both DI and DI+NMES had no effect on the aerobic performance of the ankle plantar flexors (Fig. 2B).

### Markers of skeletal muscle membrane damage

CK activity after 4 days of DI was 2.5 time lower (*p* = 0.03) than at the baseline (41 IU/l and 109 IU/l, respectively), meanwhile myoglobin level showed no changes (22.9 mg/l and 26.2 mg/l, respectively). DI+NMES significantly increased skeletal muscle membrane damage: CK activity after 4 days of DI was 2 time greater (*p* = 0.01) than at baseline (347 IU/l and 166 IU/l, respectively), as well as the myoglobin level (*p* = 0.002; 64.6 mg/l and 39.1 mg/l, respectively).

### Muscle fiber cross-sectional area and mitochondrial function in the vastus lateralis and soleus muscles

The average proportion of type I and II muscle fibers in VL before DI was 43.4(32.7-46.9)% and 56.6(53.1-67.3)%, respectively, and in Sol – 73.0(65.0-86.6)% and 21.5(10.4-25.4)%, respectively. Before DI+NMES, the average proportion of type I and II muscle fibers in VL was 43.5(41.7-45.7)% and 61.9(60.4-63.7)%, respectively, and in Sol – 79.9(74.4-88.7)% and 24.7(20.7-30.9)%, respectively. The myofiber distribution did not change significantly in both experiments. After DI, a tendency to decrease in CSA was observed in type II fibers in VL (by - 8.5(-23 – 1)%, *p* = 0.08); no changes were found in type I fibers in VL and in both fiber types in Sol (Fig. 2C). DI+NMES had no effect on both fiber types in VL and type I fibers in Sol, but reduced CSA in type II fibers in Sol (by -15(-29 – -1)%, *p* = 0.01; Fig. 2C, which is consistent with the loss of MVC in the ankle plantar flexors; Fig. 2A). Interestingly, the magnitude of change in this measure after DI differed (*p* = 0.04) from that after DI+NMES, underlining the negative effect of NMES on Sol.

DI reduced maximum ADP-stimulated mitochondrial respiration in permeabilized VL and Sol muscle fibers by -20(-33 – -5 % and -17(-25 – -7)%, respectively (*p*<0.03). In contrast, DI+NMES increased this measure by 42(3 – 90)% and 65(10 – 86)% (*p*<0.04), respectively (Fig. 2D). Moreover, in both muscles, the magnitude of change in these parameters after DI differed (*p*<0.01) from that after DI+NMES, underlining a strong positive effect of NMES on both muscles (Fig. 2D).

### Proteomic profile in the vastus lateralis and soleus muscles

A total 1890 proteins were detected in both muscles: mainly highly abundant myofibrillar, mitochondrial, cytoskeletal, sarcomeric, and ribosomal proteins, as well as heat shock and sarcoplasmic reticulum proteins (Supporting information Table S1). In VL, the relative content of 724 proteins was lower than in Sol (Fig. 3A); the majority of them (399 proteins) enriched functional categories related to mitochondrial proteins and regulators of fatty acid and carbohydrate metabolism (Fig. 3B, Supporting information Table S1 and S2). The relative content of 64 proteins was higher in VL than in Sol; these proteins were associated with calcium signaling (14 proteins), sarcomere (12 proteins), glycolysis (12 proteins), and glycogen metabolism (4 proteins) (Fig. 3B, Supporting information Table S1 and S2). In VL, the relative content of the “fast” myosin heavy chains 2X and 2A, encoded by the *MYH1* and *MYH2* genes, was 1.7 and 1.8 times higher (*q*<0.01) than in Sol, respectively, meanwhile the “slow” myosin heavy chain (*MYH7* gene) was 1.4 times less (*q*<0.01) (Supporting information Table S1). The marked difference in the proteomic profiles of VL and Sol is consistent with differences in protein expression between type I and type II muscle fibers showed in the human VL (Murgia et al. 2017; Murgia et al. 2021), as well as with differences in the transcriptomic profiles of these muscles and in the maximum rate of mitochondrial respiration in the baseline (Fig. 3A, C, and D and see below).

Further analysis showed that dry immersion had no effect on the relative content of detected proteins of either muscle. The results of the Western blot analysis for five mitochondrial proteins related to various respiratory complexes fully confirmed our proteomic data: *i*) at baseline the relative content of all mitochondrial proteins was greater in Sol than in VL, *ii*) dry immersion DI did not affect the relative content of virtually all of these proteins in either muscle (Supporting Information Fig. S4). It should be noted that, because of the high dynamic range of the muscle proteome (Deshmukh et al. 2015), the proteomic profile in this study largely did not include low abundant regulatory/signaling proteins. These proteins usually have a short half-life (Schwanhausser et al. 2011; Fornasiero et al. 2018) and thereby potentially could significantly change their relative content after 6 days of disuse.

### Transcriptomic profile in the vastus lateralis and soleus muscles

After the removal of low-expressed genes, ∼10,000 mRNAs were detected (Supporting information Table S1). As expected, in VL, compared with Sol, down-regulated mRNAs (1133) enriched terms related to aerobic and fatty acid metabolism enzymes, contractile apparatus, cell adhesion, and ion transporters, and up-regulated mRNAs (1326) were associated with regulators of translation, glycolytic enzymes, transmembrane transporters, and contractile proteins (Fig. 3C, Supporting information Table S1 and S2).

After both DI and DI+NMES, changes in gene expression in Sol (∼1600 mRNAs) were more pronounced (2.5–4 times) than in VL (∼400 mRNAs) (Fig. 4A and B, Supporting information Table S3). Interestingly, in the “mixed” fiber type composition VL, the number of genes that changed expression after DI+NMES was lower than that after DI (442 mRNAs and 607 mRNAs, respectively). In contrast, in the “slow” Sol, the response to DI+NMES was greater compared to DI (1791 mRNAs and 1527 mRNAs, respectively, Fig. 4A and C). Moreover, NMES significantly altered the transcriptomic response to immersion, inducing the expression of multiple genes that showed no overlap with genes that responded to DI (263 and 1049 mRNAs in VL and Sol, respectively) (Fig. 4C).

**Fig. 4.**
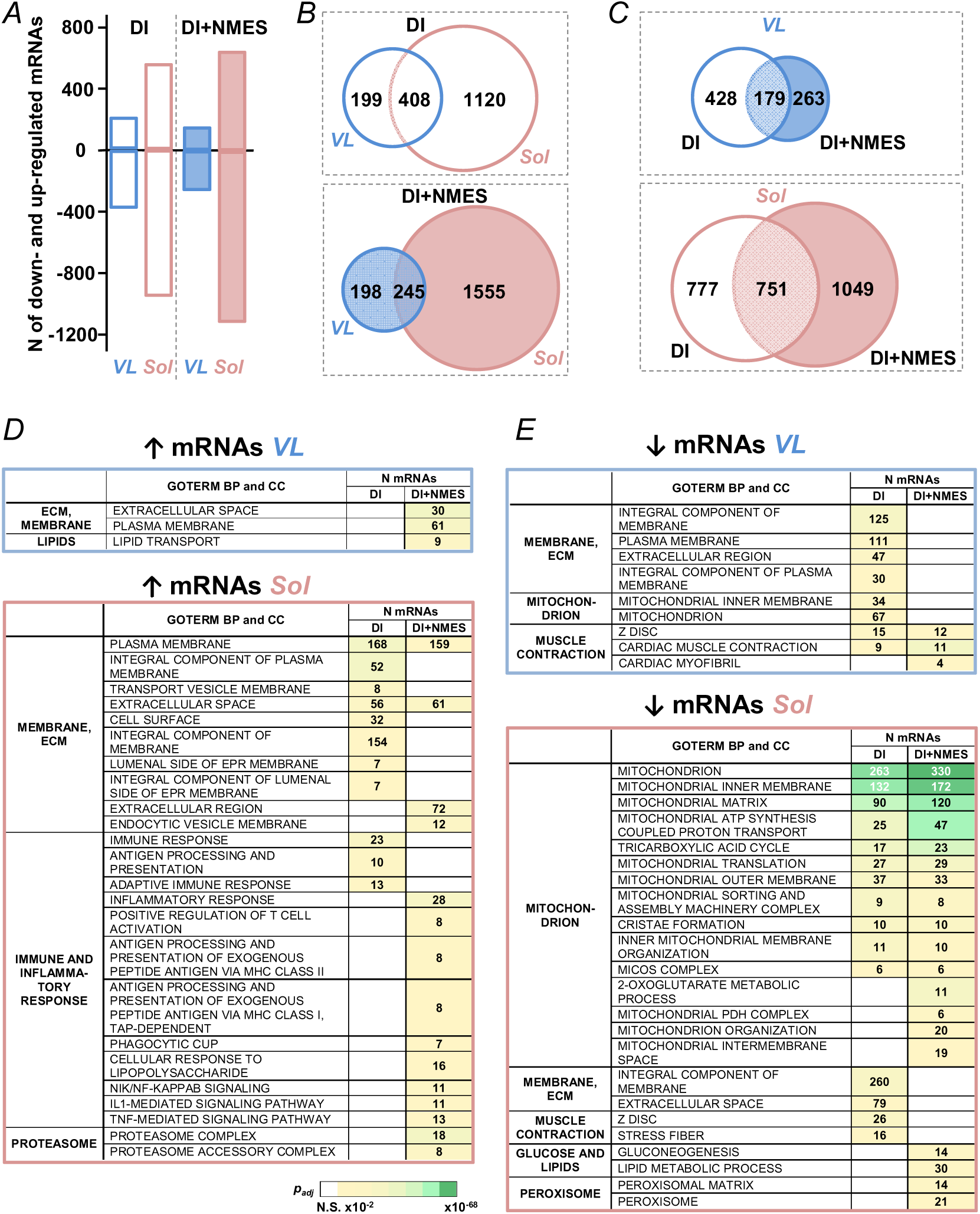
Electrical stimulation differently affected the transcriptomic response to dry immersion in the “mixed” vastus lateralis (VL) and “slow” soleus (Sol) muscles. A – The number of mRNA that changed expression in dry immersion without (DI) and with neuromuscular electrical stimulation (DI+NMES); (*n* = 10 subjects in each group). All detected protein-coding genes are presented in Supporting information Table S3. B – Transcriptomic response to DI (and DI+NMES) had little overlap between VL and Sol and was significantly more pronounced in Sol. The number of DEGs is presented. C – NMES significantly altered the transcriptomic response to immersion in both muscles. D and E – Functions of up- (D) and down-regulated (E) genes in DI and DI+NMES (all results of enrichment analysis are presented in Supporting information Table S4). The number of genes in each category is indicated, the heat map shows *p_adj_*. *N.S.* – non significant.

The small number of up-regulated genes in VL after DI (215 mRNAs) were not enriched in any functional categories (Fig. 4A and D, Supporting information Table S4). In contrast, after DI+NMES, up-regulated genes in VL (165 mRNAs) were associated with secreted proteins (30 mRNAs), plasma membrane proteins (61 mRNAs), and lipid transport (9 mRNAs) (Fig. 4A and D, Supporting information Table S4). In Sol, DI caused a large increase in gene expression (567 mRNAs) that was associated with numerous plasma membrane proteins (168 mRNAs), as well as extracellular (56 mRNAs) and immune response proteins (23 mRNAs) (Fig. 4A and D, Supporting information Table S4). Interestingly, in Sol after DI+NMES, the number of up-regulated genes increased (to 667 mRNAs); these genes enriched functional categories associated with the extracellular region (72 mRNAs), inflammatory response (28 mRNAs), and proteasome (18 mRNAs) (Fig. 4A and D, Supporting information Table S4). In addition, several genes encoding NF-kappa B inflammatory pathway proteins, cyto/myokines, and their receptors were also up-regulated (Fig. 4D and 5, Supporting information Fig. S5).

**Fig. 5.**
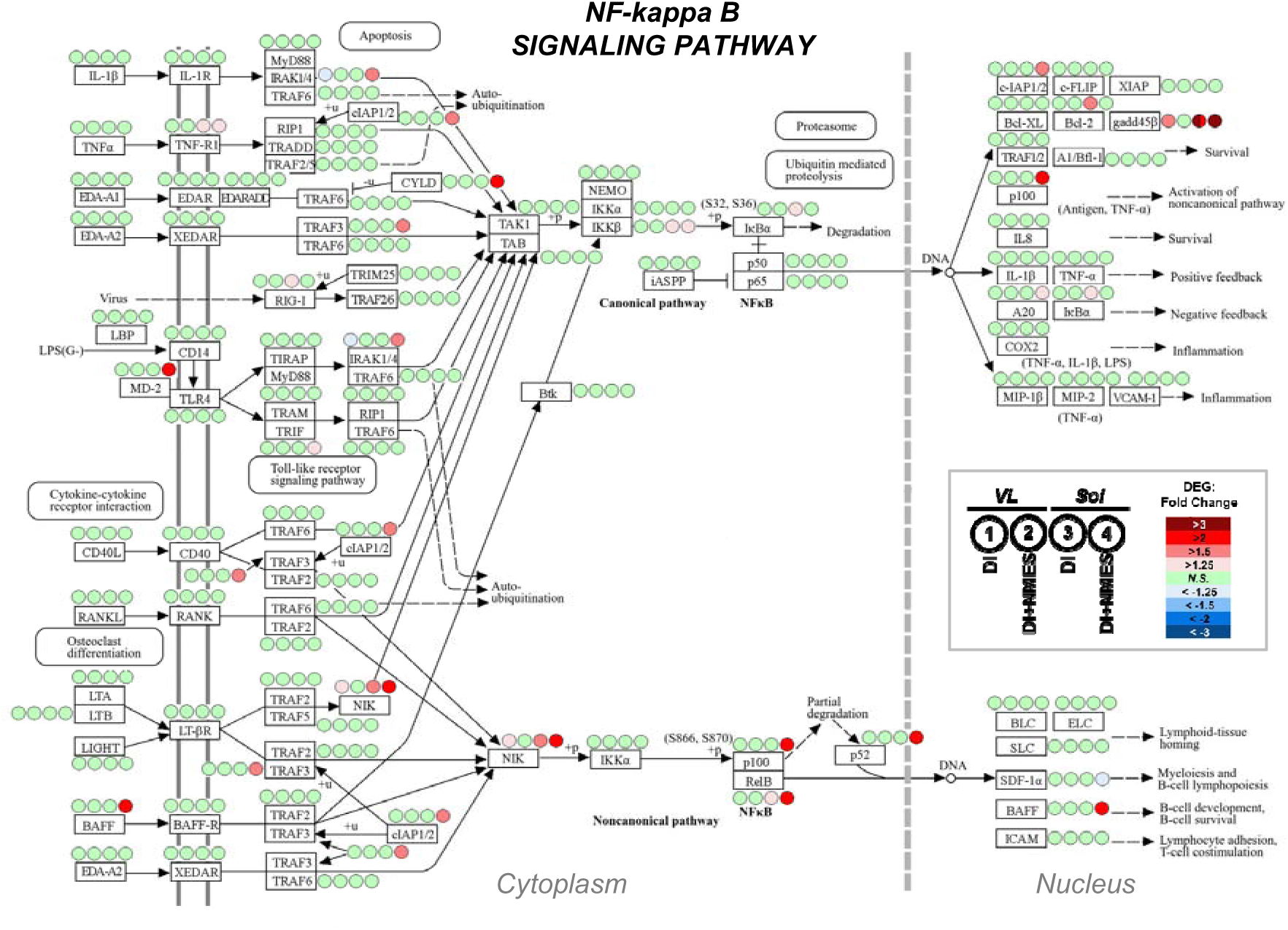
Application of neuromuscular electrical stimulation during dry immersion induced expression of mRNAs encoding proteins of the NF-kappa B inflammatory pathway in “slow” soleus muscle (Sol), but not in “mixed” vastus lateralis muscle (VL) (also see Supporting Information Fig S5). DI and DI+NMES – dry immersion without and with neuromuscular electrical stimulation, respectively; (*n* = 10 subjects in each group). Four circles next to each protein represent the change in expression of corresponding mRNA in VL and Sol after DI and DI+NMES (see the legend). The color of the circles shows the magnitude of changes in gene expression in these muscle after DI and DI+NMES (according to the heat map in the legend); *N.S.* – non-significant change.

After DI, a number of down-regulated genes in VL (392 mRNAs) were associated with a variety of membrane (125 mRNAs), mitochondrial (67 mRNAs), and sarcomeric (15 mRNAs) proteins (Fig. 4A and E, Supporting information Table S3). NMES largely prevented these negative changes: down-regulated genes were associated only with sarcomeric proteins (12 mRNAs) (Fig. 4E, Supporting information Table S4). In contrast, in Sol, DI caused the down-regulation of many genes (961 mRNAs) associated with various mitochondrial (283 mRNAs), membrane (260 mRNAs), and sarcomeric proteins (26 mRNAs) (Fig. 4A and E, Supporting information Table S4). Unexpectedly, DI+NMES increased the number of down-regulated genes in Sol (to 1133 mRNAs), especially genes encoding mitochondrial proteins (to 330 mRNAs: predominantly genes related to mitochondrial ATP synthesis coupled proton transport, tricarboxylic acid cycle, mitochondrial organization, and mitochondrial PDH complex) (Fig. 4A and E, Supporting information Table S4). Moreover, a more pronounced response (in terms of magnitude) was found for mRNAs encoding mitochondrial proteins, in particular, for the inner mitochondrial membrane (Supporting information Fig. S6). Additionally, in Sol, NMES induced a decrease in the expression of genes encoding regulators of fat metabolism (30 mRNAs) and peroxisomal proteins (21 mRNAs) (Fig. 4E). The generalized effects of DI and DI+NMES are presented in Fig. 6.

**Fig. 6.**
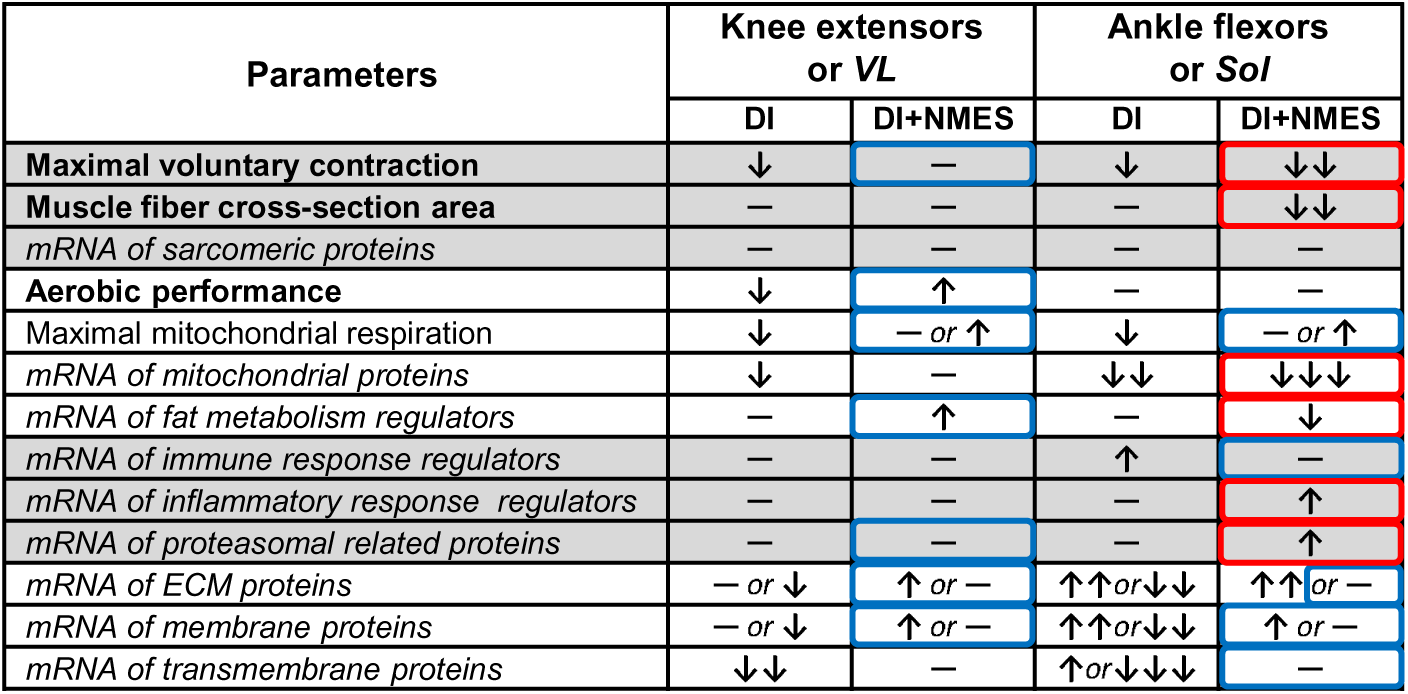
Generalized effects of dry immersion without (DI) and with low-intensity neuromuscular electrical stimulation (DI+NMES) on the functional capacity of the knee extensors/vastus lateralis muscle (VL) and the ankle plantar flexors/soleus muscle (Sol). Arrows show the direction of change; the number of arrows reflects the severity of the change/the number of genes changed expression. Blue and red boxes indicate the positive and negative effects of NMES, respectively.

## Discussion

To assess the possibility of preventing the negative effects of disuse, the effects of a week-long dry immersion without and with low-intensity NMES on the functional capacity and gene expression in the thigh and calf muscles were compared for the first time.

### NMES prevented the decline in knee extensor strength but accelerated that in ankle plantar flexors

In this study, the decline in MVC after DI was comparable to or slightly less than in the literature (Marusic et al. 2021) and did not differ between the knee extensors and ankle plantar flexors (Figs. 2 and 6). The proteomic profiles in VL and Sol were markedly different at baseline: the relative content of 788 proteins (out of 1890 detected proteins) differed between these muscles (Fig. 3). However, after 6 days of DI, no changes in the relative content of highly abundant muscle proteins, including sarcomeric proteins (constituting a half of the muscle protein mass), were found in either muscle. This may be explained by a similar decrease in the rate of synthesis of highly abundant muscle proteins and/or an increase in the rate of their degradation, as well as the fact that the expected decrease in muscle mass during weekly disuse is only a few percent (∼2.4–3.0%) (Narici and de Boer 2011; Hackney and Ploutz-Snyder 2012; Marusic et al. 2021); meanwhile, the half-life of many highly abundant proteins (sarcomeric, cytoskeletal, extracellular matrix and mitochondrial proteins, and histones) is more than a week (Fornasiero et al. 2018; Rolfs et al. 2021). The absence of changes in the expression of highly abundant contractile (as well as mitochondrial) proteins is consistent with work examining the effects of 10 days of bedrest in individual VL fibers (>200) (Murgia et al. 2022). More than two hundred proteins were found to be up- or down-regulated (out of >7500 detected); down-regulated proteins predominantly regulated the interaction of the fiber with the extracellular matrix and force transmission. It should be noted that in this work for estimating relative protein content, a label-free approach with imputation of missing data was used, and the sample size was set by the number of fibers analyzed (rather than the number of samples/volunteers), which may explain the large differences in the number of detected and differentially expressed proteins with our study.

Low-intensity NMES had different effects on different muscle groups: it prevented the disuse-induced decrease of MVC in the knee extensors, but induced the loss of fiber cross-sectional area in Sol accelerating the decrease of MVC in the ankle plantar flexors (Figs. 2 and 6). Low-intensity low- or high-frequency NMES (5%– 30% MVC, 25–50 Hz, 10–60 min/day, 5–7/week, 3–8 weeks) was previously shown to increase the muscle protein synthesis rate, size, and MVC of “mixed” knee extensor muscles in various patients with decreased muscle mass (Gibson et al. 1988; Gibson et al. 1989; Quittan et al. 2001; Harris et al. 2003) and in healthy volunteers (Stefanovska and Vodovnik 1985). An increase in MVC of the knee extensors (Lai et al. 1988) and ankle plantar flexors (Bouguetoch et al. 2021) was also shown after short-term (2–3 weeks) low-intensity high-frequency NMES (20%–25% MVC, 50–80 Hz, 5/week,) in healthy volunteers; this increase is likely due not only to the direct effect of the NMES on the muscle, but also to the neural adaptation. It can be hypothesized that a combination of the effects of dry immersion and low-intensity NMES may cause specific negative effects on Sol. NMES (including low-intensity stimulation) generates unaccustomed muscle fiber activation patterns (a synchronous and random activation of both slow and fast motor units) and primary recruit muscle volume located beneath and close to the stimulation electrodes that may lead to muscle damage and loss of MVC (Foure and Gondin 2021). In our study, electrically evoked contractions were performed with short ankle flexor muscle lengths (see Methods), which should minimize muscle damage (Foure and Gondin 2021). However, after 4 days of DI+NMES, in contrast to DI, a significant increase in the serum CK activity and myoglobin level – markers of muscle membrane damage, were found. We speculate, this effect may be particularly pronounced for Sol due to the marked decrease in its tone during dry immersion (Kozlovskaya et al. 2007). On the other hand, the negative effect of a combination of NMES and dry immersion on “slow” Sol may be related to the fact that energy supply, proteomic profile, regulation of protein synthesis and breakdown (Fig. 3) (van Wessel et al. 2010; Ciciliot et al. 2013; Murgia et al. 2017; Murgia et al. 2021), and some molecular responses to disuse (e.g., activation of inflammatory response genes (see below); Fig. 5 and Supporting information Fig. S5) differ significantly between “slow” and “fast” muscle fibers/muscles.

### NMES prevented the decline in aerobic performance only in the knee extensors

The lack of a pronounced decrease in aerobic performance in both muscle groups (Figs. 2B and 6) and change in the relative content of more than three hundred detected mitochondrial proteins in both muscles after 6 days of DI (Supporting information Table S1 and S2) is in line with studies showing no changes in time to failure/maximum power in the one-legged knee extension test after 7-10-day bedrest (Ringholm et al. 2011; Salvadego et al. 2016), as well as activity/content of mitochondrial enzymes/mitochondrial density in VL after 5-10-day bedrest (Mikines et al. 1991; Salvadego et al. 2016; Zuccarelli et al. 2021; Marshall et al. 2022; Eggelbusch et al. 2024) and 5-11-day space flight (Edgerton et al. 1995). Taken together, these data and other studies suggest that pronounced reductions in markers of mitochondrial density (for review, see (Gram et al. 2014)) and aerobic performance in endurance exercise tests that involve small muscle mass (Salvadego et al. 2011) should occur in a later stage of disuse.

Interestingly, low-intensity NMES had a small positive effect on the aerobic performance of the knee extensors only (Figs. 2B and 6) that may be related to a slightly increase in muscle oxidative capacity. Indeed, it has been shown that muscle oxidative capacity in VL of healthy volunteers can be significantly increased after just 1 week of aerobic cycling training (2 h/day) (Spina et al. 1996). Since the NMES-induced increase in aerobic performance correlates with the total stimulation time (Smart et al. 2013), it can be hypothesized that the small effect of low-intensity stimulation on the thigh muscles is due to its insufficient duration. Meanwhile the lack of effects for the calf/“slow” Sol muscles may be related to the high oxidative capacity of this muscle (and lower ability to respond to a training stimulus (Molmen et al. 2024)) and/or to activation of inflammatory response genes induced by combined exposure to immersion and NMES (Fig. 5, Supporting information Fig. S5 and see below).

DI caused a decrease in the maximum ADP-stimulated mitochondrial respiration rate in permeabilized fibers, which is consistent with several studies examining the effects of 6–10 days of disuse (Dirks et al. 2016; Dirks et al. 2020; Standley et al. 2020), but contradicts others (Salvadego et al. 2016; Edwards et al. 2020; Zuccarelli et al. 2021; Eggelbusch et al. 2024). Moreover, DI+NMES induced an opposite change in the maximal ADP-stimulated respiration, which also occurred without any significant changes in the relative content of mitochondrial proteins and aerobic performance in both muscles/muscle groups. It can be suggested that change in the maximal ADP-stimulated respiration reflects the regulation of respiration (due to post-translational modifications of respiratory enzymes, etc.) rather than mitochondrial density and protein concentration. This is indirectly confirmed by the fact that changes in the maximal ADP-stimulated respiration may occur very rapidly without changes in respiratory enzyme levels: after just three days of leg immobilization in VL (Miotto et al. 2019) and of dry immersion in Sol (Popov et al. 2023).

### NMES has differential effects on the transcriptomic response to dry immersion in the soleus and vastus lateralis muscles

Unlike mass spectrometry-based proteomic analysis, RNA-seq makes it possible to detect all expressed protein-coding genes. In line with studies showing greater negative effect of various disuse models on the phenotype of Sol than other leg muscles (LeBlanc et al. 1992; Kozlovskaya et al. 2007; Fitts et al. 2010; Tomilovskaya et al. 2019; Casuso et al. 2024), DI caused significantly greater transcriptome changes in Sol than in VL (Fig. 4). The more pronounced response in Sol and the direction of these changes are consistent with studies examining the effects of long-term (60 days) bedrest in human VL and Sol (Chopard et al. 2009) and 6 weeks of spaceflight in various leg muscles of mice (da Silveira et al. 2020). A comparison of the transcriptomic response in different muscles suggests that a longer period of disuse may lead to a significant decrease in the relative content of certain highly abundant proteins (mitochondrial, sarcomeric, etc.) in Sol that may partially explain greater negative effect of disuse on the phenotype of Sol than VL and other leg muscles.

NMES significantly altered the transcriptomic response in both muscles, but the direction of these changes differed between VL and Sol. In VL, NMES reduced most of the negative transcriptomic changes caused by immersion and, moreover, increased the expression of a number of genes encoding fat metabolism regulators and extracellular matrix proteins (Figs. 4 and 6, Supporting information Table S4), which is in line with transcriptomic changes caused by regular aerobic exercise (Popov et al. 2019; Makhnovskii et al. 2021). In contrast, in Sol, DI+NMES increased the number of down-regulated genes (genes encoding mitochondrial proteins, proteasomes, and fat metabolism regulators) and/or the magnitude of their response (Fig. 4D and E and Supporting information Fig. S6). Moreover, in Sol, DI+NMES induced an increase in genes encoding NF-kappa B pathway proteins, a number of cyto/myokines, and their receptors (Fig. 4D and 5, Supporting information Fig. S5) indicating an inflammatory response. Such response was previously shown in “slow” Sol already on the first days of disuse (Popov et al. 2023), in contrast to VL (Abadi et al. 2009). The obtained data suggest that the combination of DI and low-intensity NMES may amplify this effect in Sol, which appears to be related to both stimulation-induced muscle damage and intrinsic features of “slow” fibers in Sol. Activation of inflammatory signaling pathways has a negative impact on protein synthesis and breakdown, mitochondrial biogenesis, and other cellular processes (Costamagna et al. 2015; Thoma and Lightfoot 2018; Abu Shelbayeh et al. 2023). It can be hypothesized that the activation of inflammatory genes in Sol partially explains the negative effects of NMES after a week of dry immersion: an increase in the number of down-regulated genes encoding mitochondrial proteins (and magnitude of their response) and a decrease in the muscle fiber cross-sectional area in Sol, as well as a lack of a preventive effect on aerobic capacity and a negative impact on MVC in the ankle plantar flexors.

## Limitations

In our study, low-intensity NMES was used, therefore, mainly superficial part of the muscles was recruited. This may partly explain the variability in data obtained from the analysis of biopsy samples (CSA, mitochondrial respiration, proteomic and transcriptomic data). The variability of data after DI+NMES was also partly due to the subjective choice of stimulation intensity. However, it should be noted that this approach was chosen because it is well suited for the practical application of low-intensity NMES in patients.

The number of muscle fibers examined in our histological study was small due to the limited amount of muscle tissue available for histological assessment; this is particularly important for fast fibers, proportion of which in Sol is small. This, as well as the small sample size for some of histological analyses (*n* = 6 - 10), limits the interpretation of the data on changes in muscle fiber CSA. Additionally, because the sample size in the study was small, non-parametric statistical tests were used; therefore, some of our results may be underpowered.

An intense acute aerobic exercise was shown to induces changes in gene expression in skeletal muscle with a peak at 3 h of recovery (in terms of gene number) (Makhnovskii et al. 2021). We cannot exclude a potential acute effect of the low-intensity NMES session that preceded the second biopsy on the transcriptomic data. It should be noted that the recovery time from this NMES session to the second biopsy was 16 h, indicating that the acute effect of NMES was apparently weak. On the other hand, the combination of disuse and regular NMES sessions had opposing effects on the Sol and VL transcriptome, emphasizing the specificity of the response in these muscles rather than the acute effect of the NMES session that preceded the second biopsy. Additionally, it should be noted that gene expression changes obtained by RNA-seq were not validated by real-time PCR, which may somewhat limit the interpretation of our data.

Using the quantitative mass spectrometry-based proteomics, no changes in the relative content of highly abundant muscle proteins were observed after DI. Therefore, this approach was not used to investigate changes in the muscle proteome following DI+NMES, which somewhat limits the interpretation of our finding. Instead, a Western blot analysis was performed, which revealed no changes in proteins associated with five respiratory complexes after DI+NMES (like after DI).

## Conclusions

The present work was the first to investigate *i*) the effect of a week-long dry immersion on the phenotype and gene expression in leg muscles with significantly different functional capacities (in particular, mitochondrial respiration, proteomic and transcriptomic profile), and *ii*) the efficiency of low-intensity (∼10% MVC) NMES to prevent the negative effects of dry immersion. Dry immersion-induced decreases in MVC in both muscles were comparable and occurred without a decrease in the relative content of contractile proteins, which appears to be due to disruption of neuromuscular mechanisms rather than a decrease in muscle mass. The decrease in maximum ADP-stimulated mitochondrial respiration was also comparable in both muscles and occurred without a decrease in the relative content of mitochondrial proteins/respiratory enzymes indicating dysregulation of intrinsic respiration. At the same time, it was found that changes in the transcriptome were significantly more pronounced (2.5–4 times, in terms of number of differentially expressed genes) in the “slow” soleus muscle than in the vastus lateralis muscle. This suggests that a longer period of disuse will lead to a change (primarily a decrease) in the relative content of a number of highly abundant proteins in Sol and an accelerated decline in the functional capacity of the ankle plantar flexors.

Application of NMES during dry immersion prevented a decrease in MVC and a slight reduction in aerobic performance of the knee extensors, as well as a decrease in maximal mitochondrial respiration in permeabilized fibers and changes in the expression of genes encoding mitochondrial, extracellular matrix, and membrane proteins in the vastus lateralis muscle. In contrast, for the ankle plantar flexors/soleus muscle, NMES had a positive effect only on maximal mitochondrial respiration, but accelerated the decline in the MCV/muscle fiber cross-sectional area, which appears to be related to the activation of genes regulating the inflammatory response. Our data allows to suggest that this inflammatory response is associated with stimulation-induced muscle damage and/or intrinsic features of “slow” fibers in the soleus muscle. The data obtained open up broad prospects for the use of low-intensity NMES to prevent the negative effects of disuse for “mixed” muscles, meanwhile, the optimization of the stimulation protocol is required for “slow” muscles.

## Acknowledgements

The authors acknowledge the laboratory staffs at IBMP RAS for their excellent assistance in organization of dry immersion experiments, Motanova E.S. for assistance in the experiment on mitochondrial respiration, and are grateful for the opportunity to use the mass spectrometry equipment of the “Human Proteome” Core Facility (Institute of Biomedical Chemistry, Moscow).

## Statements and Declarations

### Data availability statement

The data supporting these findings is available in the NCBI Gene Expression Omnibus (GEO) under accession number GSE271606 and GSE271607 (RNA-seq datasets) and Supporting Information Table S1 (proteomic datasets).

### Author contributions

OIO, IVR, EST, and DVP conception or design of the work. AAB, RYZ, IIP, TFV, VGZ, MAO, NEV, N.V.S., EML, KAS, ARB, PAM, G.Y.V., R.R.G., BSS, IVR, EST, and DVP acquisition, analysis or interpretation of data for the work. AAB, RYZ, EST, and DVP drafting the work or revising it critically for important intellectual content. All authors approved the final version of the manuscript and agree to be accountable for all aspects of the work in ensuring that questions related to the accuracy or integrity of any part of the work are appropriately investigated and resolved. All persons qualify for authorship are listed.

## Supporting Information

**Supporting Information Fig. S1.** Relationships between NMES-induced muscle force (relative to maximal voluntary contraction) and intensity of NMES.

Red lines show intensity of NMES used in the main experiment (that correspond to “the point of causing unpleasant sensations”) and relative force at this intensity.

**Supporting Information Fig. S2.** A maximal isometric voluntary contraction (MVC) test and a dynamic incremental ramp test till exhaustion for the ankle plantar flexors.

A – Subject position during the MVC test and incremental ramp test for the ankle plantar flexors.

B – Dynamics of force produced in five consecutive attempts during the MVC test.

C – Dynamics of external load, motion amplitude, cycle duration, and calculated power during the incremental ramp test.

**Supporting Information Fig. S3.** A maximal isometric voluntary contraction (MVC) test and a dynamic incremental ramp test till exhaustion for the knee extensors.

A – Dynamics of knee extensors force the in five consecutive attempts during the isometric MVC test at a knee joint angle of 110 degrees.

B – Subject position during the incremental ramp test for the knee extensors.

C – Dynamics of external load, knee angle amplitude, cycle duration, and calculated power during an incremental ramp test for the knee extensors.

**Supporting Information Fig. S4.** The relative content of five mitochondrial proteins related to various mitochondrial respiratory complexes in the vastus lateralis (VL) and soleus (Sol) muscles at baseline (prior to dry immersion) and their changes after dry immersion without (DI) and with neuromuscular electrical stimulation (DI+NMES).

A – Representative immunoblot.

B – Difference between VL and Sol at baseline. * – *p* <0.05 (Mann-Whitney test)

C – Changes in VL and Sol induced by DI and DI+NMES. * – *p* <0.05 (nonparametric Wilcoxon test).

**Supporting Information Fig. S5.** Expression of mRNAs encoding cyto/myokines and their receptors (A) and proteasomal proteins (B) in the “mixed” vastus lateralis (VL) and “slow” soleus (Sol) muscles after dry immersion without (DI) and with neuromuscular electrical stimulation (DI+NMES); (*n* = 10 subjects in each group). The color of the circles and the heat map show the magnitude of changes in various muscle after various intervention for each mRNA; *N.S.* – non-significant change.

Four circles next to each protein represent the change in expression of corresponding mRNA in VL and Sol after DI and DI+NMES (see the legend). The color of the circles shows the magnitude of changes in gene expression in these muscle after DI and DI+NMES (according to the heat map in the legend); N.S. – non-significant change.

**Supporting Information Fig. S6.** Number of differentially expressed genes (DEGs) related to some enriched functional categories from Fig. 4 and their changes in expression after 6 days of dry immersion without (DI) and with neuromuscular electrical stimulation (DI+NMES); (n = 10 subjects in each group).

Common for both and unique to each intervention DEGs are shown; *p*-values relate to the difference in the magnitude of changes in expression for common DEGs between DI and DI+NMES (Mann-Whitney test).

**Supporting Information Table S1.** All detected and differentially expressed proteins (DEP) and mRNAs (DEG) in the vastus lateralis muscle (VL) *vs.* soleus muscle (S) in the baseline.

**Supporting Information Table S2.** Function enrichment analysis of differentially expressed proteins and mRNAs in the vastus lateralis muscle (VL) *vs.* soleus muscle (S) in the baseline. The main terms shown in Fig. 3 are depicted in red.

**Supporting Information Table S3.** All detected proteins and mRNAs, and differentially expressed mRNAs (DEG) in the vastus lateralis (VL) and soleus (S) muscles after 6 days of dry immersion.

**Supporting Information Table S4.** Function enrichment analysis of differentially expressed mRNAs in the vastus lateralis (VL) and soleus (S) muscles after 6 days of dry immersion.

## References

Abadi A, Glover EI, Isfort RJ, Raha S, Safdar A, Yasuda N, Kaczor JJ, Melov S, Hubbard A, Qu X et al. 2009. Limb immobilization induces a coordinate down-regulation of mitochondrial and other metabolic pathways in men and women. PLoSOne 4: e6518.

Abu Shelbayeh O, Arroum T, Morris S, Busch KB. 2023. PGC-1alpha Is a Master Regulator of Mitochondrial Lifecycle and ROS Stress Response. Antioxidants (Basel) 12.

Ade CJ, Broxterman RM, Barstow TJ. 2015. VO(2max) and Microgravity Exposure: Convective versus Diffusive O(2) Transport. Med Sci Sports Exerc 47: 1351–1361.

Amirova L, Avdeeva M, Shishkin N, Gudkova A, Guekht A, Tomilovskaya E. 2022. Effect of modulated electromyostimulation on the motor system of elderly neurological patients. Pilot study of Russian currents also known as Kotz currents. Front Physiol 13: 921434.

Bouguetoch A, Martin A, Grospretre S. 2021. Insights into the combination of neuromuscular electrical stimulation and motor imagery in a training-based approach. Eur J Appl Physiol 121: 941–955.

Casuso RA, Huertas JR, Aragon-Vela J. 2024. The role of muscle disuse in muscular and cardiovascular fitness: A systematic review and meta-regression. Eur J Sport Sci 24: 812–823.

Chopard A, Lecunff M, Danger R, Lamirault G, Bihouee A, Teusan R, Jasmin BJ, Marini JF, Leger JJ. 2009. Large-scale mRNA analysis of female skeletal muscles during 60 days of bed rest with and without exercise or dietary protein supplementation as countermeasures. Physiol Genomics 38: 291–302.

Ciciliot S, Rossi AC, Dyar KA, Blaauw B, Schiaffino S. 2013. Muscle type and fiber type specificity in muscle wasting. Int J Biochem Cell Biol 45: 2191–2199.

Costamagna D, Costelli P, Sampaolesi M, Penna F. 2015. Role of Inflammation in Muscle Homeostasis and Myogenesis. Mediators Inflamm 2015: 805172.

Cresswell AG, Loscher WN, Thorstensson A. 1995. Influence of gastrocnemius muscle length on triceps surae torque development and electromyographic activity in man. Exp Brain Res 105: 283–290.

da Silveira WA, Fazelinia H, Rosenthal SB, Laiakis EC, Kim MS, Meydan C, Kidane Y, Rathi KS, Smith SM, Stear B et al. 2020. Comprehensive Multi-omics Analysis Reveals Mitochondrial Stress as a Central Biological Hub for Spaceflight Impact. Cell 183: 1185–1201 e1120.

Deshmukh AS, Murgia M, Nagaraj N, Treebak JT, Cox J, Mann M. 2015. Deep proteomics of mouse skeletal muscle enables quantitation of protein isoforms, metabolic pathways, and transcription factors. MolCell Proteomics 14: 841–853.

Dirks ML, Miotto PM, Goossens GH, Senden JM, Petrick HL, van Kranenburg J, van Loon LJC, Holloway GP. 2020. Short-term bed rest-induced insulin resistance cannot be explained by increased mitochondrial H2 O2 emission. J Physiol 598: 123–137.

Dirks ML, Wall BT, van de Valk B, Holloway TM, Holloway GP, Chabowski A, Goossens GH, van Loon LJ. 2016. One Week of Bed Rest Leads to Substantial Muscle Atrophy and Induces Whole-Body Insulin Resistance in the Absence of Skeletal Muscle Lipid Accumulation. Diabetes 65: 2862–2875.

Edgerton VR, Zhou MY, Ohira Y, Klitgaard H, Jiang B, Bell G, Harris B, Saltin B, Gollnick PD, Roy RR et al. 1995. Human fiber size and enzymatic properties after 5 and 11 days of spaceflight. J Appl Physiol (1985) 78: 1733–1739.

Edwards SJ, Smeuninx B, McKendry J, Nishimura Y, Luo D, Marshall RN, Perkins M, Ramsay J, Joanisse S, Philp A et al. 2020. High-dose leucine supplementation does not prevent muscle atrophy or strength loss over 7 days of immobilization in healthy young males. Am J Clin Nutr 112: 1368–1381.

Eggelbusch M, Charlton BT, Bosutti A, Ganse B, Giakoumaki I, Grootemaat AE, Hendrickse PW, Jaspers Y, Kemp S, Kerkhoff TJ et al. 2024. The impact of bed rest on human skeletal muscle metabolism. Cell Rep Med 5: 101372.

Fitts RH, Trappe SW, Costill DL, Gallagher PM, Creer AC, Colloton PA, Peters JR, Romatowski JG, Bain JL, Riley DA. 2010. Prolonged space flight-induced alterations in the structure and function of human skeletal muscle fibres. J Physiol 588: 3567–3592.

Fornasiero EF, Mandad S, Wildhagen H, Alevra M, Rammner B, Keihani S, Opazo F, Urban I, Ischebeck T, Sakib MS et al. 2018. Precisely measured protein lifetimes in the mouse brain reveal differences across tissues and subcellular fractions. Nat Commun 9: 4230.

Foure A, Gondin J. 2021. Skeletal Muscle Damage Produced by Electrically Evoked Muscle Contractions. Exerc Sport Sci Rev 49: 59–65.

Gibson JN, Morrison WL, Scrimgeour CM, Smith K, Stoward PJ, Rennie MJ. 1989. Effects of therapeutic percutaneous electrical stimulation of atrophic human quadriceps on muscle composition, protein synthesis and contractile properties. Eur J Clin Invest 19: 206–212.

Gibson JN, Smith K, Rennie MJ. 1988. Prevention of disuse muscle atrophy by means of electrical stimulation: maintenance of protein synthesis. Lancet 2: 767–770.

Gram M, Dahl R, Dela F. 2014. Physical inactivity and muscle oxidative capacity in humans. Eur J Sport Sci 14: 376–383.

Hackney KJ, Ploutz-Snyder LL. 2012. Unilateral lower limb suspension: integrative physiological knowledge from the past 20 years (1991-2011). Eur J Appl Physiol 112: 9–22.

Harris S, LeMaitre JP, Mackenzie G, Fox KA, Denvir MA. 2003. A randomised study of home-based electrical stimulation of the legs and conventional bicycle exercise training for patients with chronic heart failure. Eur Heart J 24: 871–878.

Karavidas A, Arapi SM, Pyrgakis V, Adamopoulos S. 2010. Functional electrical stimulation of lower limbs in patients with chronic heart failure. Heart Fail Rev 15: 563–579.

Koryak YA. 2020. Isokinetic Force and Work Capacity After Long-Duration Space Station Mir and Short-Term International Space Station Missions. Aerosp Med Hum Perform 91: 422–431.

Kozlovskaya IB, Sayenko IV, Sayenko DG, Miller TF, Khusnutdinova DR, Melnik KA. 2007. Role of support afferentation in control of the tonic muscle activity. Acta Astronautica 60: 285–294.

Kurochkina NS, Orlova MA, Vigovskiy MA, Zgoda VG, Vepkhvadze TF, Vavilov NE, Makhnovskii PA, Grigorieva OA, Boroday YR, Philippov VV et al. 2024. Age-related changes in human skeletal muscle transcriptome and proteome are more affected by chronic inflammation and physical inactivity than primary aging. Aging Cell doi:10.1111/acel.14098: e14098.

Lai HS, Domenico GD, Strauss GR. 1988. The effect of different electro-motor stimulation training intensities on strength improvement. Aust J Physiother 34: 151–164.

LeBlanc AD, Schneider VS, Evans HJ, Pientok C, Rowe R, Spector E. 1992. Regional changes in muscle mass following 17 weeks of bed rest. J Appl Physiol (1985) 73: 2172–2178.

Maffiuletti NA, Green DA, Vaz MA, Dirks ML. 2019. Neuromuscular Electrical Stimulation as a Potential Countermeasure for Skeletal Muscle Atrophy and Weakness During Human Spaceflight. Front Physiol 10: 1031.

Makhnovskii PA, Bokov RO, Kolpakov FA, Popov DV. 2021. Transcriptomic Signatures and Upstream Regulation in Human Skeletal Muscle Adapted to Disuse and Aerobic Exercise. Int J Mol Sci 22.

Marshall RN, Smeuninx B, Seabright AP, Morgan PT, Atherton PJ, Philp A, Breen L. 2022. No effect of five days of bed rest or short-term resistance exercise prehabilitation on markers of skeletal muscle mitochondrial content and dynamics in older adults. Physiol Rep 10: e15345.

Marusic U, Narici M, Simunic B, Pisot R, Ritzmann R. 2021. Nonuniform loss of muscle strength and atrophy during bed rest: a systematic review. J Appl Physiol (1985) 131: 194–206.

Mayr W, Bijak M, Girsch W, Hofer C, Lanmuller H, Rafolt D, Rakos M, Sauermann S, Schmutterer C, Schnetz G et al. 1999. MYOSTIM-FES to prevent muscle atrophy in microgravity and bed rest: preliminary report. Artif Organs 23: 428–431.

Mikines KJ, Richter EA, Dela F, Galbo H. 1991. Seven days of bed rest decrease insulin action on glucose uptake in leg and whole body. J Appl Physiol (1985) 70: 1245–1254.

Miotto PM, McGlory C, Bahniwal R, Kamal M, Phillips SM, Holloway GP. 2019. Supplementation with dietary omega-3 mitigates immobilization-induced reductions in skeletal muscle mitochondrial respiration in young women. FASEB J 33: 8232–8240.

Molmen KS, Almquist NW, Skattebo O. 2024. Effects of Exercise Training on Mitochondrial and Capillary Growth in Human Skeletal Muscle: A Systematic Review and Meta-Regression. Sports Med doi:10.1007/s40279-024-02120-2.

Murgia M, Ciciliot S, Nagaraj N, Reggiani C, Schiaffino S, Franchi MV, Pisot R, Simunic B, Toniolo L, Blaauw B et al. 2022. Signatures of muscle disuse in spaceflight and bed rest revealed by single muscle fiber proteomics. PNAS Nexus 1: pgac086.

Murgia M, Nogara L, Baraldo M, Reggiani C, Mann M, Schiaffino S. 2021. Protein profile of fiber types in human skeletal muscle: a single-fiber proteomics study. Skelet Muscle 11: 24.

Murgia M, Toniolo L, Nagaraj N, Ciciliot S, Vindigni V, Schiaffino S, Reggiani C, Mann M. 2017. Single Muscle Fiber Proteomics Reveals Fiber-Type-Specific Features of Human Muscle Aging. Cell Rep 19: 2396–2409.

Narici MV, de Boer MD. 2011. Disuse of the musculo-skeletal system in space and on earth. Eur J Appl Physiol 111: 403–420.

Niess F, Fiedler GB, Schmid AI, Laistler E, Frass-Kriegl R, Wolzt M, Moser E, Meyerspeer M. 2018. Dynamic multivoxel-localized (31) P MRS during plantar flexion exercise with variable knee angle. NMR Biomed 31: e3905.

Nuhr MJ, Pette D, Berger R, Quittan M, Crevenna R, Huelsman M, Wiesinger GF, Moser P, Fialka-Moser V, Pacher R. 2004. Beneficial effects of chronic low-frequency stimulation of thigh muscles in patients with advanced chronic heart failure. Eur Heart J 25: 136–143.

Poltavskaya MG, Sviridenko VP, Brand AV, Andreev DE, Koryak YA, Veliev GO, Dikur ON, Kulikov VM, Vaisman YD, Tomilovskaya ES. 2021. The Use of “Space” Electrical Myostimulation in Clinical Cardiology on Earth. Hum Physiology 47: 382–390.

Popov DV, Lysenko EA, Bokov RO, Volodina MA, Kurochkina NS, Makhnovskii PA, Vyssokikh MY, Vinogradova OL. 2018. Effect of aerobic training on baseline expression of signaling and respiratory proteins in human skeletal muscle. Physiological Reports 6: e13868.

Popov DV, Makhnovskii PA, Shagimardanova EI, Gazizova GR, Lysenko EA, Gusev OA, Vinogradova OL. 2019. Contractile activity-specific transcriptome response to acute endurance exercise and training in human skeletal muscle. Am J Physiol Endocrinol Metab 316: E605–E614.

Popov DV, Makhnovskii PA, Zgoda VG, Gazizova GR, Vepkhvadze TF, Lednev EM, Motanova ES, Lysenko EA, Orlov OI, Tomilovskaya ES. 2023. Rapid changes in transcriptomic profile and mitochondrial function in human soleus muscle after 3-day dry immersion. J Appl Physiol (1985) 134: 1256–1264.

Quittan M, Wiesinger GF, Sturm B, Puig S, Mayr W, Sochor A, Paternostro T, Resch KL, Pacher R, Fialka-Moser V. 2001. Improvement of thigh muscles by neuromuscular electrical stimulation in patients with refractory heart failure: a single-blind, randomized, controlled trial. Am J Phys Med Rehabil 80: 206–214; quiz 215-206, 224.

Ried-Larsen M, Aarts HM, Joyner MJ. 2017. Effects of strict prolonged bed rest on cardiorespiratory fitness: systematic review and meta-analysis. J Appl Physiol (1985) 123: 790–799.

Ringholm S, Bienso RS, Kiilerich K, Guadalupe-Grau A, Aachmann-Andersen NJ, Saltin B, Plomgaard P, Lundby C, Wojtaszewski JF, Calbet JA et al. 2011. Bed rest reduces metabolic protein content and abolishes exercise-induced mRNA responses in human skeletal muscle. Am J Physiol Endocrinol Metab 301: E649–658.

Rolfs Z, Frey BL, Shi X, Kawai Y, Smith LM, Welham NV. 2021. An atlas of protein turnover rates in mouse tissues. Nat Commun 12: 6778.

Salvadego D, Keramidas ME, Brocca L, Domenis R, Mavelli I, Rittweger J, Eiken O, Mekjavic IB, Grassi B. 2016. Separate and combined effects of a 10-d exposure to hypoxia and inactivity on oxidative function in vivo and mitochondrial respiration ex vivo in humans. J Appl Physiol (1985) 121: 154–163.

Salvadego D, Keramidas ME, Kolegard R, Brocca L, Lazzer S, Mavelli I, Rittweger J, Eiken O, Mekjavic IB, Grassi B. 2018. PlanHab(*) : hypoxia does not worsen the impairment of skeletal muscle oxidative function induced by bed rest alone. J Physiol 596: 3341–3355.

Salvadego D, Lazzer S, Marzorati M, Porcelli S, Rejc E, Simunic B, Pisot R, di Prampero PE, Grassi B. 2011. Functional impairment of skeletal muscle oxidative metabolism during knee extension exercise after bed rest. J Appl Physiol (1985) 111: 1719–1726.

Schwanhausser B, Busse D, Li N, Dittmar G, Schuchhardt J, Wolf J, Chen W, Selbach M. 2011. Global quantification of mammalian gene expression control. Nature 473: 337–342.

Shiba N, Matsuse H, Takano Y, Yoshimitsu K, Omoto M, Hashida R, Tagawa Y, Inada T, Yamada S, Ohshima H. 2015. Electrically Stimulated Antagonist Muscle Contraction Increased Muscle Mass and Bone Mineral Density of One Astronaut - Initial Verification on the International Space Station. PLoS One 10: e0134736.

Smart NA, Dieberg G, Giallauria F. 2013. Functional electrical stimulation for chronic heart failure: a meta-analysis. Int J Cardiol 167: 80–86.

Spina RJ, Chi MM, Hopkins MG, Nemeth PM, Lowry OH, Holloszy JO. 1996. Mitochondrial enzymes increase in muscle in response to 7-10 days of cycle exercise. JApplPhysiol 80: 2250–2254.

Standley RA, Distefano G, Trevino MB, Chen E, Narain NR, Greenwood B, Kondakci G, Tolstikov VV, Kiebish MA, Yu G et al. 2020. Skeletal Muscle Energetics and Mitochondrial Function Are Impaired Following 10 Days of Bed Rest in Older Adults. J GerontolA BiolSciMedSci 75: 1744–1753.

Stefanovska A, Vodovnik L. 1985. Change in muscle force following electrical stimulation. Dependence on stimulation waveform and frequency. Scand J Rehabil Med 17: 141–146.

Thoma A, Lightfoot AP. 2018. NF-kB and Inflammatory Cytokine Signalling: Role in Skeletal Muscle Atrophy. Adv Exp Med Biol 1088: 267–279.

Tomilovskaya E, Shigueva T, Sayenko D, Rukavishnikov I, Kozlovskaya I. 2019. Dry Immersion as a Ground-Based Model of Microgravity Physiological Effects. Front Physiol 10: 284.

van Wessel T, de Haan A, van der Laarse WJ, Jaspers RT. 2010. The muscle fiber type-fiber size paradox: hypertrophy or oxidative metabolism? EurJApplPhysiol 110: 665–694.

Vikne H, Strom V, Pripp AH, Gjovaag T. 2020. Human skeletal muscle fiber type percentage and area after reduced muscle use: A systematic review and meta-analysis. Scand J Med Sci Sports 30: 1298–1317.

Ward AR, Shkuratova N. 2002. Russian Electrical Stimulation: The Early Experiments. Physical Therapy 82: 1019–1030.

Yarmanova EN, Kozlovskaya IB, Khimoroda NN, Fomina EV, Reeves JM. 2015. Evolution of russian microgravity countermeasures. Aerospace Medicine and Human Performance 86: A32–A37.

Yu SH, Kyriakidou P, Cox J. 2020. Isobaric Matching between Runs and Novel PSM-Level Normalization in MaxQuant Strongly Improve Reporter Ion-Based Quantification. J Proteome Res 19: 3945–3954.

Zanotti E, Felicetti G, Maini M, Fracchia C. 2003. Peripheral muscle strength training in bed-bound patients with COPD receiving mechanical ventilation: effect of electrical stimulation. Chest 124: 292–296.

Zuccarelli L, Baldassarre G, Magnesa B, Degano C, Comelli M, Gasparini M, Manferdelli G, Marzorati M, Mavelli I, Pilotto A et al. 2021. Peripheral impairments of oxidative metabolism after a 10-day bed rest are upstream of mitochondrial respiration. J Physiol 599: 4813–4829.

