## Supplemental Figure S1 for "Multidirectional effect of low-intensity neuromuscular electrical stimulation on gene expression and phenotype in thigh and calf muscles after one week of disuse"

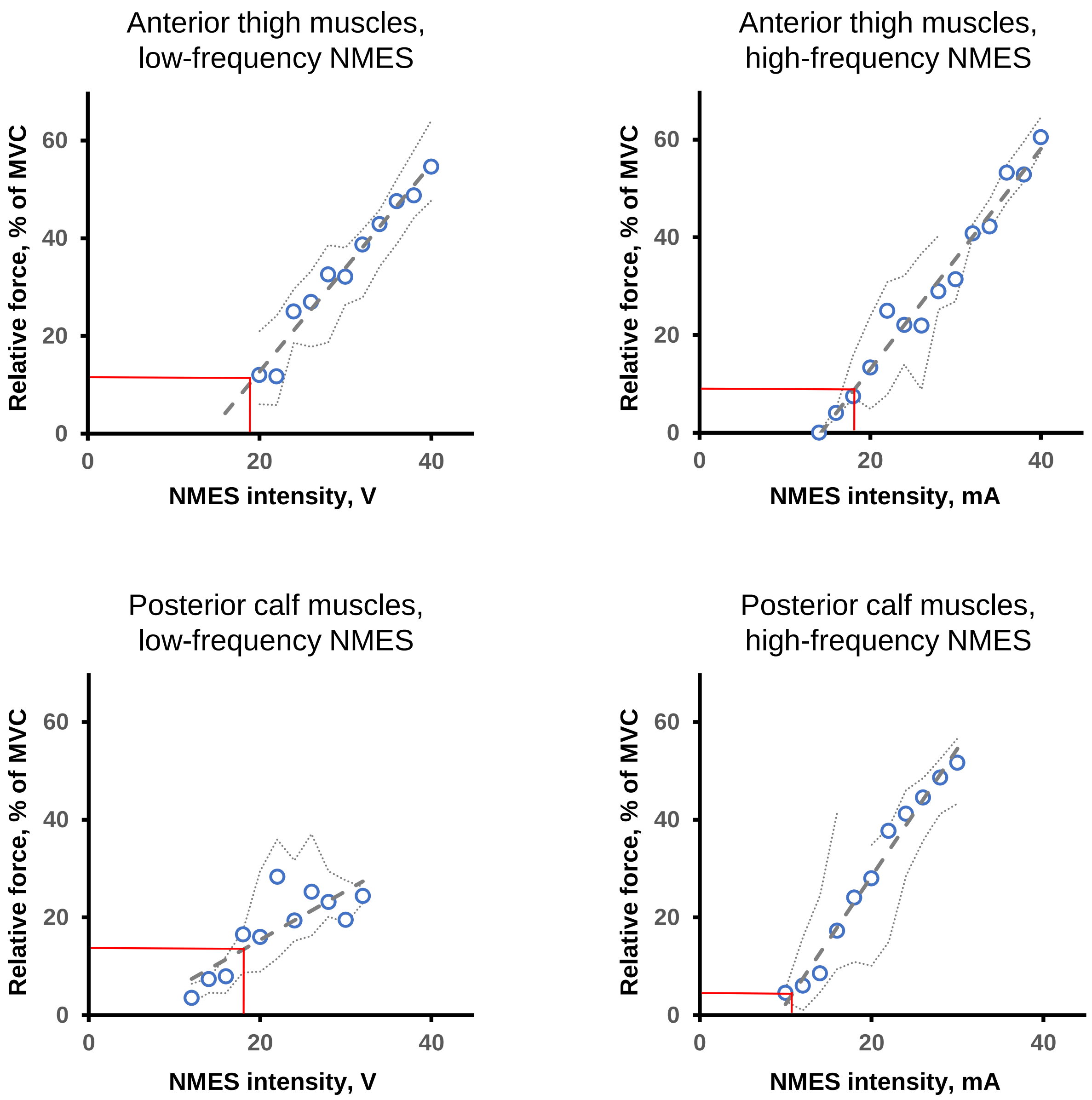


**Supporting Information Fig. S1.** Relationships between NMES-induced muscle force (relative to maximal voluntary contraction) and intensity of NMES.

Red lines show intensity of NMES used in the main experiment (that correspond to "the point of causing unpleasant sensations") and relative force at this intensity.

.
