## Supplemental Figure S2 for "Multidirectional effect of low-intensity neuromuscular electrical stimulation on gene expression and phenotype in thigh and calf muscles after one week of disuse"

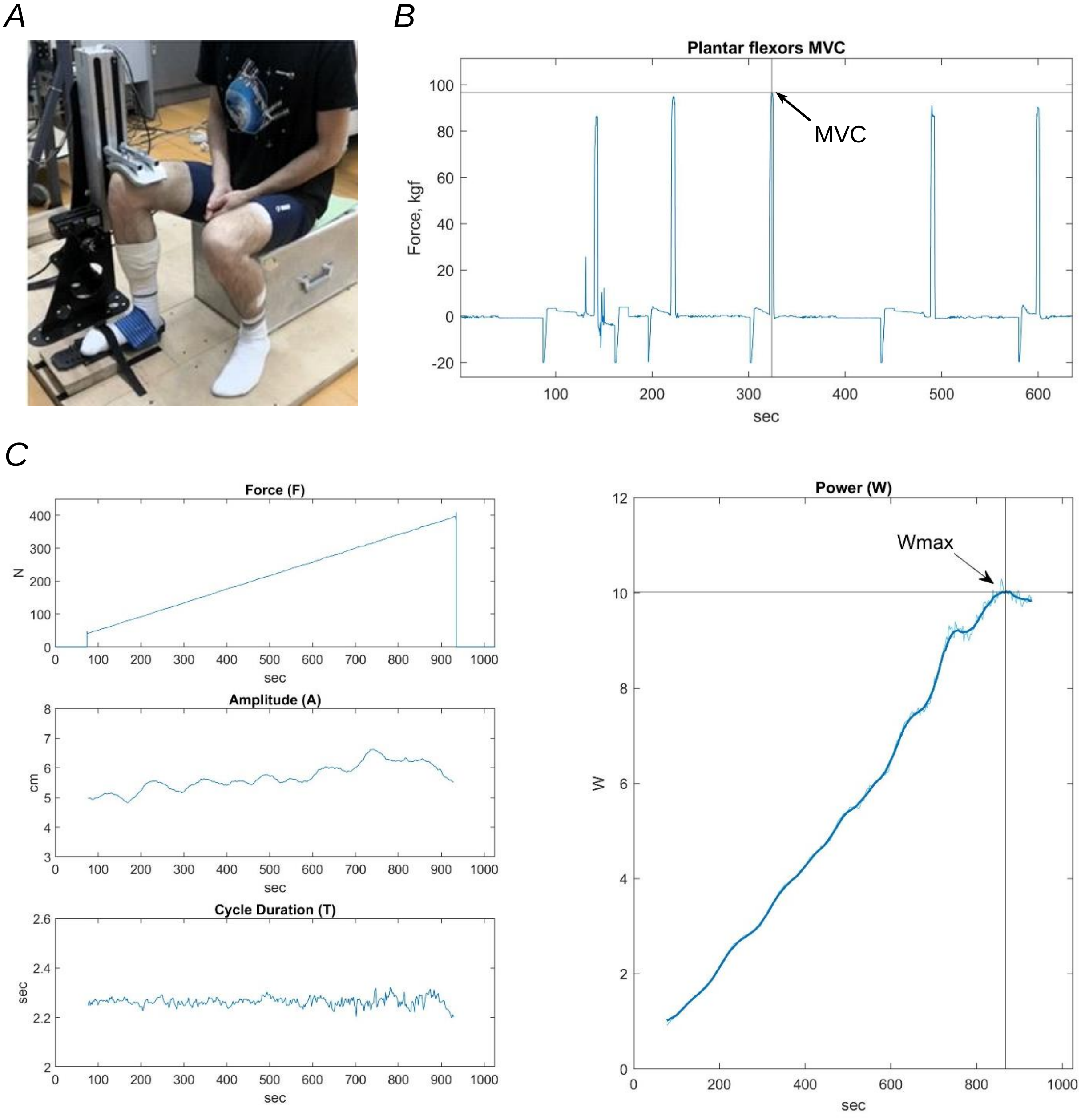


**Supporting Information Fig. S2.** A maximal isometric voluntary contraction (MVC) test and a dynamic incremental ramp test till exhaustion for the ankle plantar flexors.

A – Subject position during the MVC test and incremental ramp test for the ankle plantar flexors.

B – Dynamics of force produced in five consecutive attempts during the MVC test.

C – Dynamics of external load, motion amplitude, cycle duration, and calculated power during the incremental ramp test.
