## Supplemental Figure S3 for "Multidirectional effect of low-intensity neuromuscular electrical stimulation on gene expression and phenotype in thigh and calf muscles after one week of disuse"

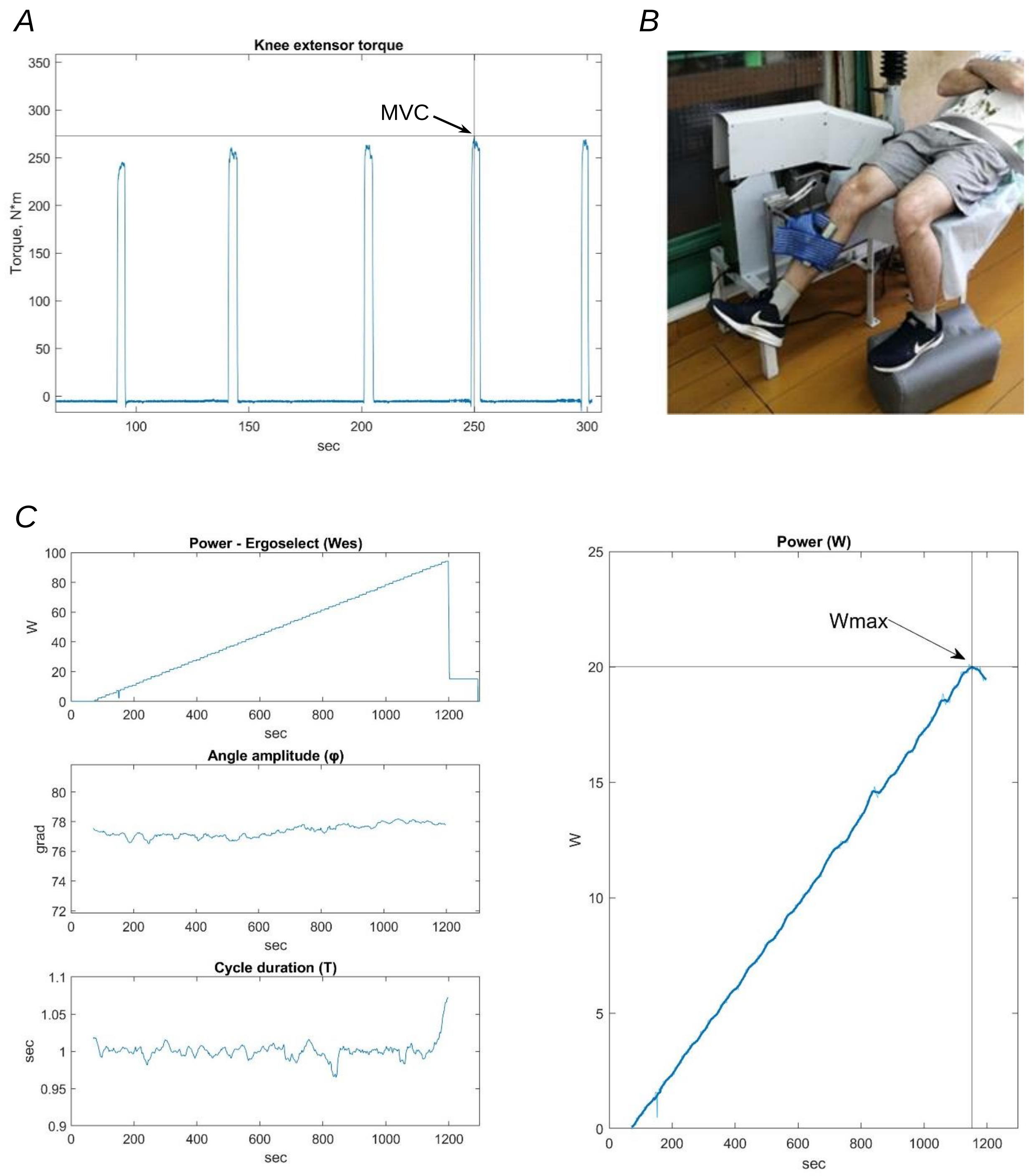


**Supporting Information Fig. S3.** A maximal isometric voluntary contraction (MVC) test and a dynamic incremental ramp test till exhaustion for the knee extensors.

A – Dynamics of knee extensors force the in five consecutive attempts during the isometric MVC test at a knee joint angle of 110 degrees.

B – Subject position during the incremental ramp test for the knee extensors.

C – Dynamics of external load, knee angle amplitude, cycle duration, and calculated power during an incremental ramp test for the knee extensors.
