## Supplemental Figure S4 for "Multidirectional effect of low-intensity neuromuscular electrical stimulation on gene expression and phenotype in thigh and calf muscles after one week of disuse"

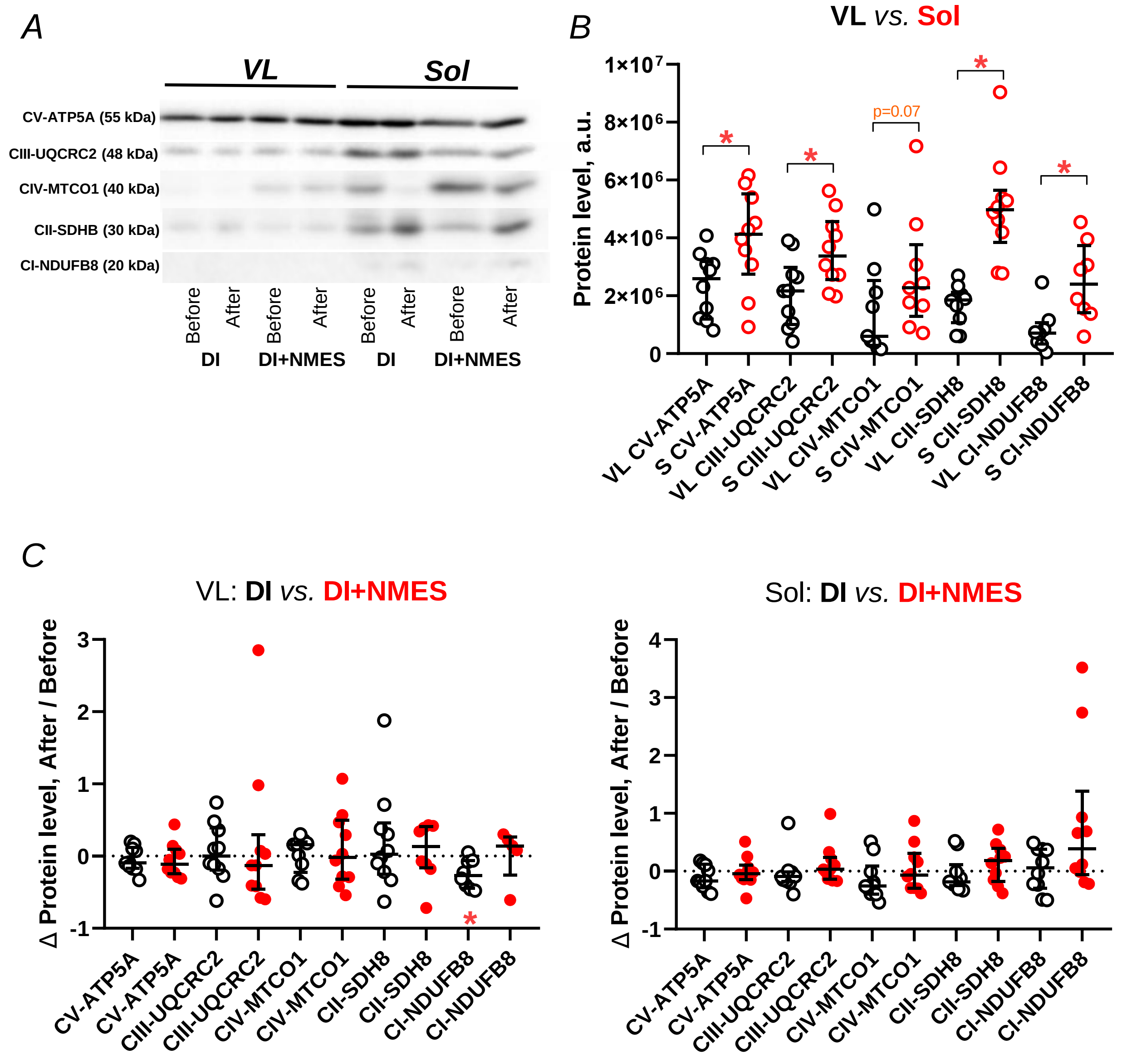


**Supporting Information Fig. S4.** The relative content of five mitochondrial proteins related to various mitochondrial respiratory complexes in the vastus lateralis (VL) and soleus (Sol) muscles at baseline (prior to dry immersion) and their changes after dry immersion without (DI) and with neuromuscular electrical stimulation (DI+NMES).

A – Representative immunoblot.

B – Difference between VL and Sol at baseline. * – p <0.05 (Mann-Whitney test)

C – Changes in VL and Sol induced by DI and DI+NMES. * – p <0.05 (nonparametric Wilcoxon test).
