## Supplemental Figure S5 for "Multidirectional effect of low-intensity neuromuscular electrical stimulation on gene expression and phenotype in thigh and calf muscles after one week of disuse"

A

### CYTOKINE-CYTOKINE RECEPTOR INTERECTION

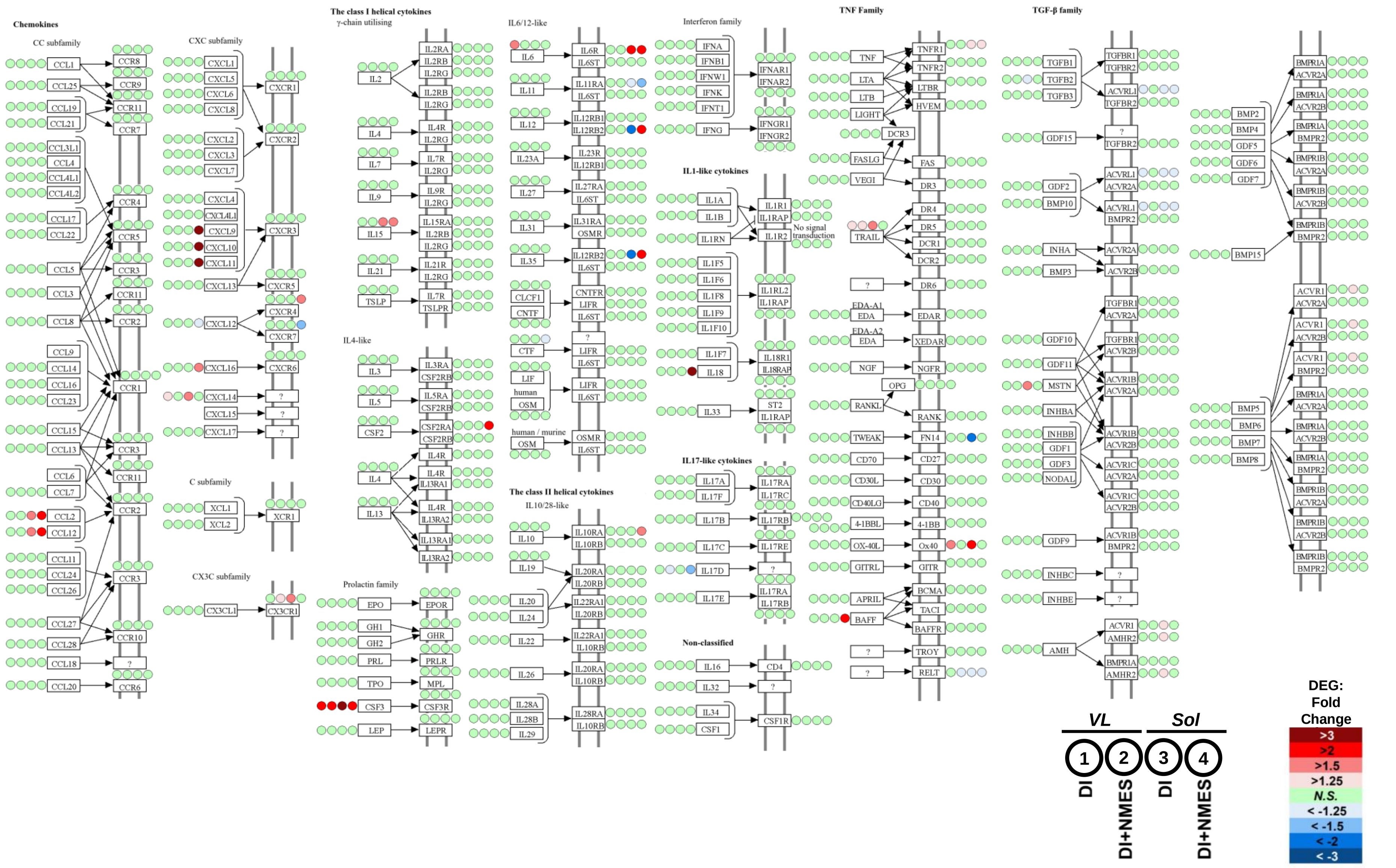

*B*

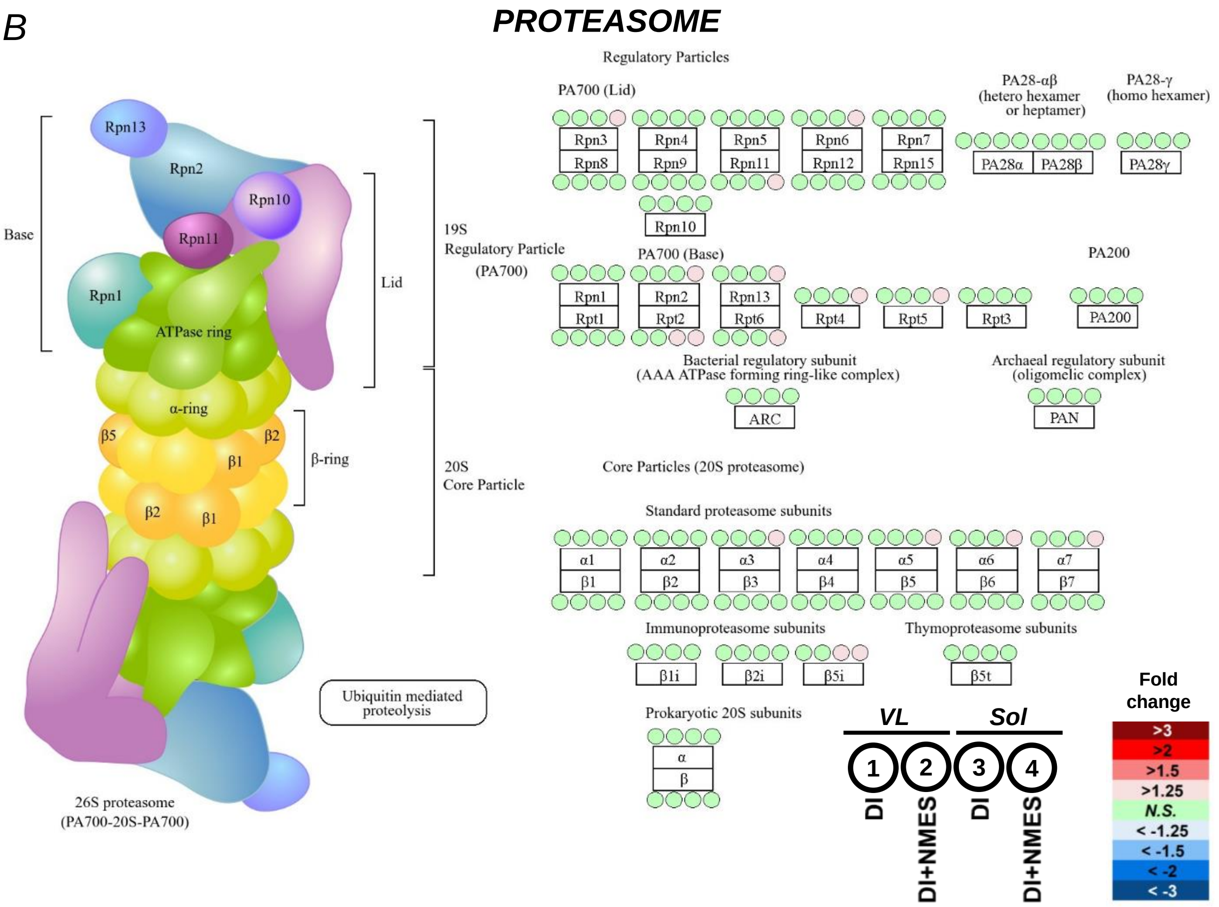


**Supporting Information Fig. S5.** Expression of mRNA encoding cyto/myokines and their receptors (A) and proteasomal proteins (B) in the "mixed" vastus lateralis (VL) and "slow" soleus (Sol) muscles after dry immersion without (DI) and with neuromuscular electrical stimulation (DI+NMES); (n = 10 subjects in each group).

Four circles next to each protein represent the change in expression of corresponding mRNA in VL and Sol after DI and DI+NMES (see the legend). The color of the circles shows the magnitude of changes in gene expression in these muscle after DI and DI+NMES (according to the heat map in the legend); N.S. – non-significant change.
