## Supplemental Figure S5 for "Multidirectional effect of low-intensity neuromuscular electrical stimulation on gene expression and phenotype in thigh and calf muscles after one week of disuse"

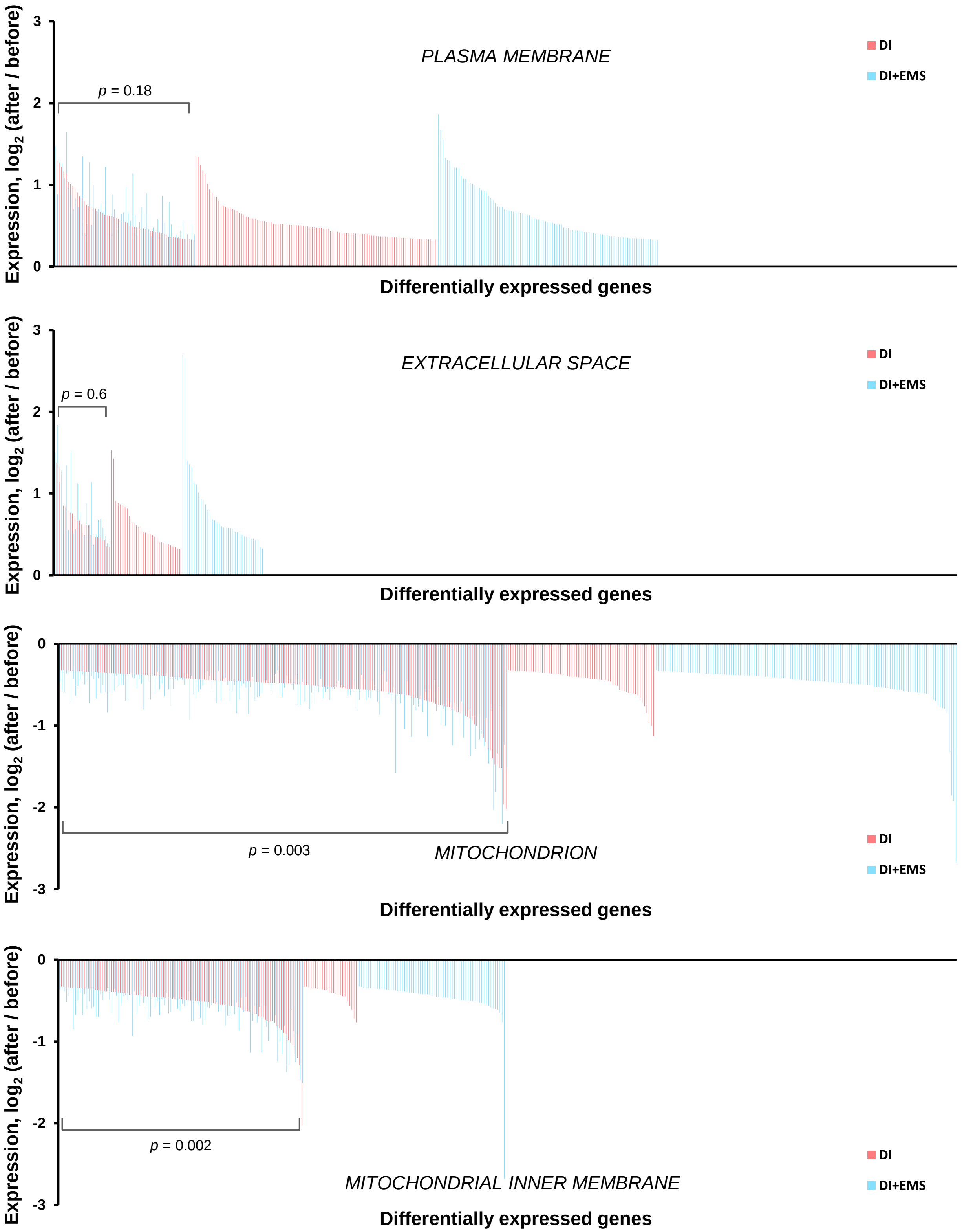


**Supporting Information Fig. S6.** Number of differentially expressed genes (DEGs) related to some enriched functional categories from Fig. 4 and their changes in expression after 6 days of dry immersion without (DI) and with neuromuscular electrical stimulation (DI+NMES); (n = 10 subjects in each group).

Common for both and unique to each intervention DEGs are shown; *p*-values relate to the difference in the magnitude of changes in expression for common DEGs between DI and DI+NMES (Mann-Whitney test).
